# The antitumor peptide CIGB-552 is active in B-cell lymphomas and enhances BTK-targeted therapies

**DOI:** 10.64898/2026.11.25.769102

**Authors:** Maidel Carpio Alvarez, Alberto J. Arribas, Luciano Cascione, Emma Pesenti, Andrea Rinaldi, Matteo Pecoraro, Filippo Spriano, Luca Aresu, Afua Adjeiwaa Mensah, Riccardo Moia, Nawar Maher, Federico Simonetta, Gianluca Gaidano, Franco Cavalli, Julio Raúl Fernández Massó, Francesco Bertoni

## Abstract

Despite advances in targeted therapies, acquired resistance remains a major challenge in B-cell lymphomas. CIGB-552 is a synthetic antitumor peptide that modulates COMMD1-associated signaling pathways, but its activity in hematological malignancies remains unexplored. We investigated CIGB-552 in marginal zone lymphoma (MZL), mantle cell lymphoma (MCL), and diffuse large B-cell lymphoma (DLBCL) models, focusing on drug-resistant MZL and combinations with BTK-targeted agents. CIGB-552 reduced viability across lymphoma cell lines, with the greatest sensitivity in MZL and MCL, and retained activity in MZL models with acquired resistance to BTK, PI3K, or BCL2 inhibitors, but not in non-neoplastic B-cells. Peptide uptake varied among models and correlated with sensitivity, whereas treatment induced context-dependent apoptotic and non-apoptotic cell death. Integrated transcriptomic and proteomic analyses in MZL cells identified reduced B-cell receptor, NF-κB, PI3K/AKT/mTOR, and MAPK signaling alongside induction of cellular stress-response programs. Immunoblotting supported the suppression of key survival proteins and showed increased COMMD1 abundance. CIGB-552 enhanced the activity of ibrutinib in parental and resistant MZL models, and in a canine DLBCL cell line, and showed favorable interactions with additional BTK inhibitors and a BTK degrader. CIGB-552 also inhibited growth in primary lymphoma samples, and its combination with ibrutinib increased activity in primary MZL cells. These preclinical findings support further investigation of CIGB-552 as a therapeutic candidate for B-cell lymphomas, particularly in combination with BTK-targeted agents.

**Key points:**

- CIGB-552 reduces viability in B-cell lymphoma models, with prominent activity in MZL and MCL and retained activity in drug-resistant MZL derivatives.
- CIGB-552 enhances BTK-targeted treatment in lymphoma models, including primary MZL cells.

## Introduction

B-cell lymphomas are characterized by dysregulated signaling pathways that support malignant B-cell survival, including B-cell receptor (BCR) and NF-κB signaling (1–3). Inhibitors of Bruton tyrosine kinase (BTK) and phosphoinositide 3-kinase (PI3K) have improved treatment options, but intrinsic and acquired resistance continues to limit their benefit (2,3). Consequently, agents with mechanisms that complement existing targeted therapies are needed, particularly for patients whose disease progresses after pathway-directed treatment (1).

Therapeutic peptides offer a distinct class of antitumor agents with properties that can differ from those of small molecules and antibodies (4–7). CIGB-552 is a synthetic peptide derived from an antimicrobial peptide and has shown antitumor activity both *in vitro* and *in vivo* in solid tumor models (8) (12). It can enter cells through energy-dependent endocytosis or direct membrane transduction, depending on experimental conditions (9).

CIGB-552 increases the abundance of copper metabolism MURR1 domain-containing protein 1 (COMMD1), a regulator of NF-κB signaling, protein trafficking, and cellular stress responses (10,11). Reported downstream effects include reduced RelA/p65 signaling, interference with hypoxia-inducible factor 1 activity, and altered superoxide dismutase 1 maturation (10,11). Whether these mechanisms contribute to the activity of CIGB-552 in lymphoma remains to be established.

Furthermore, a phase I trial in patients with recurrent, refractory, or progressive solid tumors showed that CIGB-552 is well tolerated at doses up to 4.7 mg administered three times per week, with no significant adverse effects (12).

Here, we evaluate CIGB-552 in human B-cell lymphoma cell lines and primary samples, including MZL models with acquired resistance to BTK, PI3K, and BCL2 inhibitors. We characterize cellular uptake, cell-sdeath responses, and treatment-associated molecular changes and examine combinations with BTK inhibitors and a BTK degrader.

## Materials and Methods

### Cell line culture

Established human B-cell lymphoma cell lines were used in this study, derived from marginal zone lymphoma (MZL) or sharing similar biological features (VL51, Karpas1718 (K1718), SSK41, ESKOL, HAIRM, and HC-1) with derivatives with acquired secondary resistance to targeted therapies (13–17), from mantle cell lymphoma (MCL) (MAVER1, JEKO1, REC1, SP49 and GRANTA-519), and from diffuse large B cell lymphoma (DLBCL), comprising activated B-cell–like (ABC) models (TMD8, SU-DHL-2, OCI-LY10) and germinal center B-cell–like (GCB) models (PFEIFFER, VAL, SU-DHL-16, and OCI-LY7). The canine CLBL-1 cell line was used as an additional comparative model. Primary cells from patients with MZL, MCL, relapsed/refractory MCL, and DLBCL were used as *ex vivo* models. Finally, the immortalized lymphoblastoid cell line CB33 was used as a control model of non-malignant B lymphocytes.

Cell lines were cultured in RPMI-1640 (VL51, K1718, SSK41, ESKOL, HAIRM, HC-1, SP49, PFEIFFER, MAVER1, REC1, JEKO1, TMD8, SU-DHL-2, VAL, CLBL-1, primary cells and CB33), DMEM (GRANTA-519), or IMDM (OCI-LY7, OCI-LY10, and SU-DHL-16) supplemented with 10% fetal bovine serum (FBS), 1% L- glutamine, and a penicillin-streptomycin-neomycin mixture (approximately 5,000 units penicillin, 5 mg streptomycin, and 10 mg neomycin/mL; Sigma-Aldrich). Cells were maintained at 37 °C in a humidified atmosphere containing 5% CO_2_. All cell lines were periodically tested for mycoplasma contamination using the MycoAlert Mycoplasma Detection Kit (Lonza, Visp, Switzerland).

### Treatments and cell viability assays

CIGB-552 and fluorescein-conjugated CIGB-552 (CIGB-552-Flu) were obtained from the Center for Genetic Engineering and Biotechnology (Havana, Cuba). Ibrutinib and zanubrutinib were purchased from Selleckchem (Houston, TX, USA), while pirtobrutinib and NX-5948 were obtained from MedChemExpress (Monmouth Junction, NJ, USA) and a-Tocopherol (vitamin E) were purchased from Sigma (Buchs, Switzerland).

Cells were seeded in non-tissue-culture–treated 96-well plates at the following densities: 10,000 cells per well for VL51, MAVER1, JEKO1, TMD8, SU-DHL-2, OCI-LY10 and VAL; 20,000 cells per well for SSK41, ESKOL, HAIRM, HC-1; GRANTA-519, PFEIFFER, SU-DHL-16, and OCI-LY7; and 30,000 cells per well for K1718, REC1, SP49, CLBL-1 and CB33. CIGB-552 was dissolved in RPMI-1640 medium or Milli-Ǫ water, whereas all other compounds were dissolved in dimethyl sulfoxide (DMSO). Compounds were then serially diluted in complete medium within selected concentration ranges and added to cells in triplicate. Cells were incubated for 72 hours at 37 °C and 5% CO_2_. Cells treated with 0.1% DMSO served as negative controls, and wells containing medium only were included as blanks.

Cell viability was assessed using the MTT assay. MTT (3-(4,5-dimethylthiazol-2-yl)-2,5-diphenyltetrazolium bromide; Sigma, Switzerland) was prepared as a 5 mg/mL stock solution in phosphate-buffered saline (PBS) and filter-sterilized. MTT solution (20 µL) was added to each well, followed by incubation at 37 °C for 4 hours. Cells were lysed using 25% SDS lysis buffer, and absorbance was measured at 560 nm using Cytation 3 microplate reader (BioTek, Winooski, VT, USA). The 72-hour MTT assay was performed with CIGB-552 up to 300 µM as a single agent, and in combination CIGB-552 up to 25 µM with ibrutinib up to 100 nM, Zanubrutinib at 61.7 nM, Pirtobrutinib at 100 nM and NX-5948 at 1 nM.

Drug sensitivity was evaluated by calculating IC50 values using a four-parameter logistic model on log- scaled doses (PharmacoGx R package (18)). Combination effects were assessed using the Chou–Talalay combination index (19), the MuSyC algorithm (20), and the highest single agent (HSA) and zero interaction potency (ZIP) models implemented in the SynergyFinder R package (21).

### Live-cell proliferation analysis (Incucyte assay)

For live-cell proliferation assays, 96-well plates were coated with 50 μL per well of 0.01% (w/v) poly-L- ornithine (Sigma-Aldrich, P4957) before cell seeding. Cell densities were adjusted according to the experimental model and treatment conditions as follows: 5,000 cells/well for parental and ibrutinib- resistant VL51 cells treated with CIGB-552 as a single agent; 10,000 cells/well for parental and venetoclax- resistant SSK41 cells, parental and idelalisib-resistant Karpas1718 (K1718) cells, parental and ibrutinib- resistant VL51 cells treated with the combination of CIGB-552 and ibrutinib, and CLBL-1 cells; and 20,000 cells/well for CB33 cells and primary lymphoma cells from patients with MZL, MCL, relapsed/refractory MCL (MCL-RR), and DLBCL.

Cells were treated with 0.1% DMSO as control, CIGB-552, ibrutinib, or a combination of both agents, as indicated for each experiment. In cell-line experiments, CIGB-552 was used at 0.3, 1.5, 3, or 6 μM, and ibrutinib at 5, 50 or 100 pM. Primary lymphoma cells were treated with CIGB-552 (6 μM), ibrutinib (1 μM), or the combination of both agents.

Cell proliferation was monitored for 3–6 days using the Incucyte S3 Live-Cell Imaging System (Sartorius). Image acquisition and time-course analyses were performed using Incucyte software. Growth curves were normalized to the initial measurement at time zero to facilitate comparisons across experimental conditions.

### Primary lymphoma samples

Fresh lymph node specimens were processed under sterile conditions in a biological safety cabinet. Primary samples were mechanically dissociated using sterile scalpels in Petri dishes kept on ice to obtain cell suspensions. The resulting suspensions were diluted in 0.9% NaCl and centrifuged, and the cell pellets were collected for cryopreservation. Two cryoprotective solutions were prepared: Solution A, containing 40% RPMI-1640 medium and 60% FBS, and Solution B, containing 80% RPMI-1640 medium and 20% DMSO. Cell pellets were gently resuspended in an equal mixture of Solutions A and B, resulting in a final freezing medium containing 60% RPMI-1640, 30% FBS, and 10% DMSO. Cell suspensions were aliquoted into cryovials (approximately 30 million cells per vial), initially stored at −80°C in isopropanol-based freezing containers, and subsequently transferred to liquid nitrogen for long-term preservation. Before functional analyses, thawed primary cells were characterized by flow cytometry which determined the presence of more than 39.3% B cells, using anti-CD20 and anti-CD3 antibodies (15–20 min on ice, protected from light; 10,000 events acquired per sample). Patient samples were collected and used in accordance with ethical guidelines. Primary cells were exposed to CIGB-552 at the indicated concentrations. Cell growth was monitored for 136 hours using IncuCyte live-cell imaging and quantified as confluence area and area under the curve.

### Internalization assay

For peptide internalization studies, cells were seeded in 24-well plates at a density of 150,000 cells per well in 500 µL of complete RPMI medium and incubated at 37 °C for 24 hours. Fluorescein-conjugated CIGB- 552 was added to a final concentration of 5 µM and incubated for 1 hour at either 4 °C or 37 °C (for analysis of temperature effect) or 2 hours (for internalization assay).

Following incubation, cells were washed twice with cold PBS. Surface-bound fluorescein-labeled peptide was quenched using trypan blue (0.04 mg/mL), and non-viable cells were identified using propidium iodide (PI; 5 µg/mL). Samples were analyzed within 30–60 minutes using a FACSCanto II flow cytometer (BD Biosciences, Allschwil, Switzerland). Data acquisition and analysis were performed using FlowJo software (version 10.0.7r2; BD Life Sciences, Ashland, OR, USA).

### Cell cycle and cell death analysis

Cell cycle or apoptosis assay were assessed in 1 million cells per condition in 4 mL of complete RPMI medium, treated with PBS or CIGB-552. For the cell cycle, CIGB-552 was added for 24 or 48 hours, and after treatment, cells were harvested, washed with 1%-FBS-PBS, fixed in cold ethanol 100% and stored at −20 °C for 24 hours. Then, cells were stained with PI (50 µg/mL; Sigma-Aldrich) in 1%-FBS-PBS containing RNase A (75 kU/mL; Sigma-Aldrich) and analyzed for DNA content using a FACSCanto II flow cytometer. Cell cycle phases were quantified using FlowJo software and the Watson Pragmatic model. The role of oxidative stress was assessed by pre-treating K1718 and SSK41 cells with 100 µM α-tocopherol (40 min) prior to exposure to 50 µM CIGB-552 for 48 hours, followed by cell cycle analysis.

Apoptosis was evaluated by annexin V–FITC and PI staining. Cells were treated with CIGB-552 at 2× IC50 for 48 and 72 hours, washed twice with cold 1%-FBS-PBS, and centrifuged at 1,200 rpm for 5 minutes at 4 °C. Cells were incubated with annexin V–FITC (5 µL in 200 µL binding buffer) for 10 minutes at room temperature in the dark, washed, and subsequently stained with PI (10 µL in 200 µL binding buffer). Samples were analyzed by flow cytometry, and apoptotic and necrotic populations were quantified using FlowJo software.

### Immunoblotting

Cells were seeded in T25 flasks at densities of 3 × 10⁶ cells for VL51 and 5 × 10⁶ cells for K1718 and SSK41 in 10 mL of complete RPMI medium. After treatment, cells were harvested, pelleted, and lysed using M-PER (Mammalian Protein Extraction Reagent, Thermo Scientific, 78501) supplemented with Halt Protease and Phosphatase Inhibitor Cocktail, EDTA-free (Thermo Scientific, 78440), for 30 minutes. Lysates were clarified by centrifugation at 4 °C for 30 minutes.

Protein concentration was determined using the BCA assay (Thermo Scientific, 23225), and 30 µg of total protein per sample was resolved by 4–20% SDS–PAGE and transferred to nitrocellulose membranes. Membranes were incubated overnight with primary antibodies against AKT/p(S473)-AKT, NF-κB1, NF-κB2, RelA/p(S536)-RelA, MCL1, BCL2, PARP1, COMMD1, Cyclin D1, and ERK/p(Y204)-ERK (Table S1), followed by horseradish peroxidase (HRP)-conjugated secondary antibodies (Amersham Life Science) for 1 hour. Proteins were visualized using Western Antares ECL reagent (Cyanagen, XLS142). Vinculin was used as a loading control.

### Transcriptome profiling

VL51 marginal zone lymphoma cells (approximately 2 × 10⁶ cells per sample) were treated with CIGB-552 at its IC₅₀ concentration (26 µM) for 24 h. Experiments were performed using three independent biological replicates per condition. Suspension cells were cultured in horizontally positioned flasks to maximize the available culture surface area and facilitate peptide–cell interactions. Total RNA was independently extracted using the AllPrep DNA/RNA Mini Kit (Ǫiagen, Hilden, Germany) according to the manufacturer’s instructions. RNA-seq libraries were prepared following poly(A) enrichment using the NEBNext UltraExpress RNA Library Prep Kit (New England Biolabs). Sequencing was performed on an Illumina NextSeq 2000 platform using a P2 100-cycle reagent kit with a 120–8–8 cycle configuration.

The quality of raw RNA-seq reads was assessed during the preprocessing step using FastǪC (v0.11.5) (22). Low-quality reads/bases and adaptors were removed using Trimmomatic (v0.35) (23). The resulting high- quality trimmed reads were aligned to the human reference genome (GRCh38, version 113) using STAR (24), a spliced-read aligner that supports the alignment of reads spanning multiple exons. Samples were considered of good quality if more than 85% of sequencing reads aligned to the reference genome. Gene expression quantification was performed using HTSeq-count (25) with Ensembl gene annotation for GRCh38.113 as reference. Expression values were provided in tab-delimited format. We filtered the dataset to retain genes with counts-per-million (CPM) values greater than 5 in at least half of the samples. The data were normalized using the TMM (trimmed mean of M-values) method from the edgeR package (26) and transformed to log2 CPM using the cpm() function. Differential gene expression for each comparison of interest was computed using limma, following TMM normalization and voom transformation (27). The RNA-seq datasets generated in this study have been deposited in the NCBI Gene Expression Omnibus (GEO) database under accession number GSE354291.

### Proteomics

VL51 MZL cells (approximately 2 × 10⁶ cells per sample) from three independent biological replicates were treated in horizontally positioned flasks to maximize the available culture surface area and facilitate peptide–cell interactions. Following treatment, cells were washed twice with cold PBS, snap-frozen in liquid nitrogen, and stored at −80 °C until analysis.

Proteins were extracted in 8 M urea/50 mM ammonium bicarbonate by sonication using a Bioruptor system (Diagenode), reduced with dithiothreitol, alkylated with iodoacetamide, and sequentially digested with LysC (Wako Fujifilm) and trypsin (Promega), each at a 1:100 enzyme-to-protein ratio (w/w). Peptides were purified using C18 StageTips and analyzed by LC–MS/MS on a nanoElute 2 HPLC system coupled to a timsTOF HT mass spectrometer (Bruker) via a CaptiveSpray nanoelectrospray source. Peptides were separated on an in-house-packed C18-AǪ column (ReproSil-Pur, 1.9 μm; Dr. Maisch HPLC GmbH) using a 60-min linear gradient from 2% to 35% acetonitrile in 0.1% formic acid at a flow rate of 300 nL/min. Data were acquired in data-independent acquisition (DIA)-PASEF mode.

Raw data were processed using DIA-NN v1.8.1 against a deep-learning-predicted spectral library generated from the February 2024 release of the human UniProt database. Searches used Trypsin/P specificity, allowing one missed cleavage, with N-terminal methionine excision enabled and carbamidomethylation of cysteine as a fixed modification. Precursor and fragment mass tolerances were determined automatically for each run (10–20 ppm). The resulting protein-group matrix (*report.pg_matrix.tsv*) was used for downstream analyses.

Proteomic data analysis was performed in R using the DEP package (28). Protein intensity matrices were imported from a MaxǪuant output file and protein identifiers were made unique using the make_unique function. An experimental design table was constructed to encode sample conditions and biological replicates. A SummarizedExperiment object was generated with make_se. Proteins identified in all replicates of at least one condition were retained (filter_missval, threshold = 0). Variance- stabilizing normalization was applied using normalize_vsn. Missing values were imputed using a k-nearest neighbor approach (impute, fun = “knn”), and alternative imputation strategies (MinProb and manual left- shifted Gaussian) were evaluated for quality control. Multidimensional scaling (MDS) plots were used to assess sample clustering before and after imputation, and batch effects related to replicate structure were explored using removeBatchEffect. Differential protein abundance was assessed using limma-based linear modeling (test_diff), focusing on both specific contrasts and all pairwise comparisons. Statistically significant differentially expressed proteins were identified using a false discovery rate threshold of 0.05 and log2 fold-change cutoffs (|log2FC| ≥ log2(2) or as specified). Results were visualized using volcano plots, principal component analysis, and heatmaps of significant proteins. All statistical outputs were exported for downstream analyses.

### Artificial Intelligence Generated Content

The authors used OpenAI’s ChatGPT (GPT-6, OpenAI, San Francisco, CA, USA) to assist in text preparation and drafting. ChatGPT was also used to support exploratory data mining of author-generated datasets. All content and analyses were reviewed, edited, and verified by the authors, who take full responsibility for the manuscript.

## Results

### CIGB-552 has dose-dependent antitumor activity in human and canine B-cell lymphoma cell lines

We investigated the antitumor activity of CIGB-552 in cell lines derived from MZL, including some with secondary resistance to targeted agents. We first evaluated cell viability after 72 hours of CIGB-552 treatment in a panel of six MZL cell lines (VL51, K1718, SSK41, ESKOL, HAIRM, and HC-1). CIGB-552 induced a dose-dependent reduction in viability across all models, confirming its cytotoxic activity in lymphoma cells (Figure 1A). The median IC50 across parental MZL cell lines was 18.2 µM (95% CI: 14.1– 22.3).

**Figure 1.**
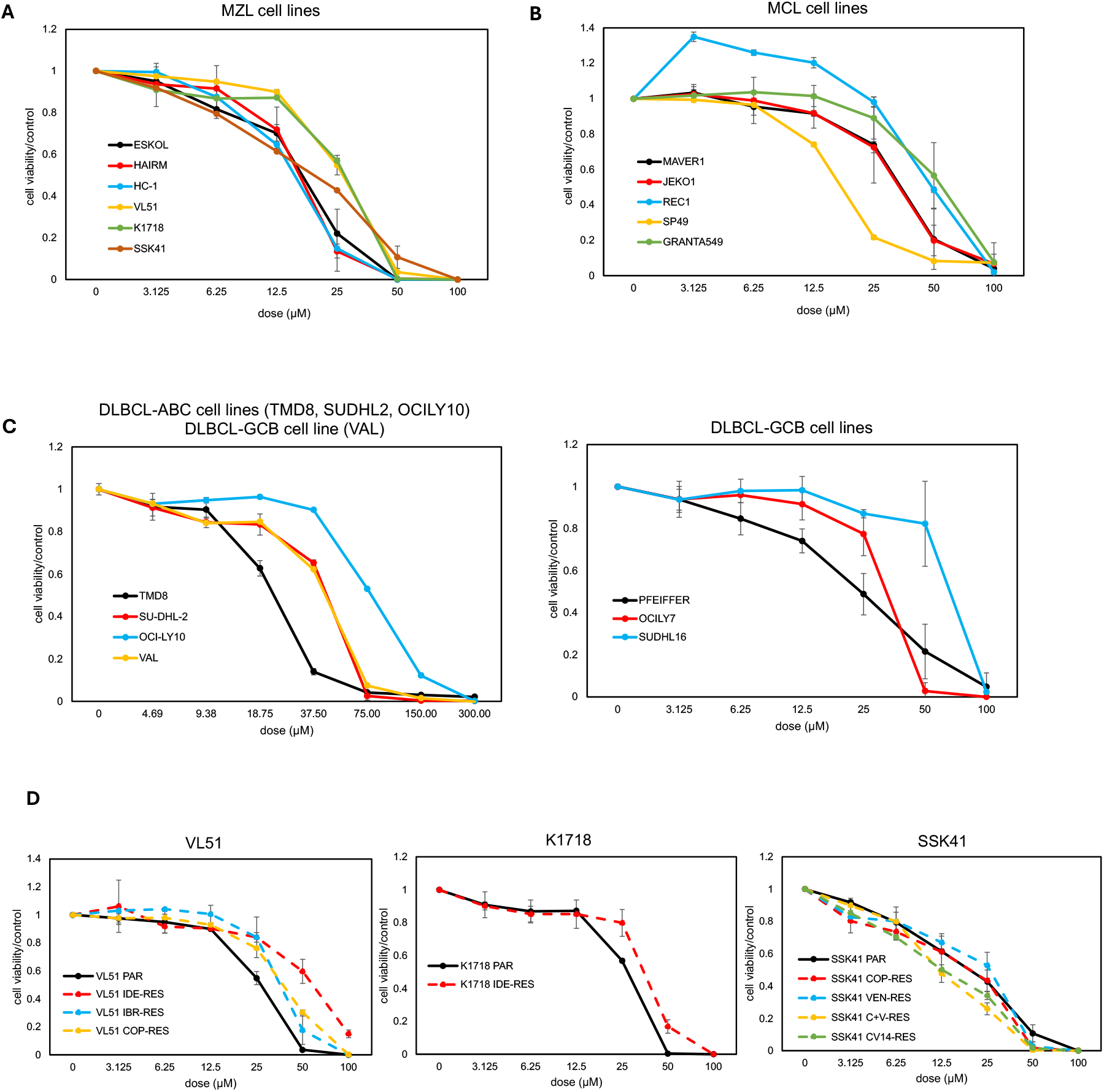
Antitumor activity of CIGB-552 in lymphoma cell lines. CIGB-552 activity was evaluated by MTT assay following 72 h of exposure in MZL (A), MCL (B), DLBCL (C) cell lines, and MZL cell lines with acquired resistance to targeted agents (D). CIGB-552 was tested using two-fold serial dilutions, starting at 100 µM for MZL, MCL, DLBCL-GCB, and drug-resistant cell lines, and at 300 µM for DLBCL-ABC cell lines and the DLBCL-GCB cell line VAL. Error bars represent the standard deviation (SD). Representative dose–response curves from at least two independent experiments are shown.

Following the activity observed in the MZL cell lines, we also tested CIGB-552 in cell lines derived from MCL (n.=5) and diffuse large B-cell lymphoma (DLBCL), either activated B-cell-like (ABC; n.=3) or germinal center B-cell (GCB; n.=4) subtypes. Cells were exposed to increasing concentrations of CIGB-552 for 72 hours. CIGB-552 induced a dose-dependent reduction in cell viability across all MCL models (Figure 1B), the calculated median IC50 was 33.9 µM (95% CI: 21.9–45.9). The peptide induced a dose-dependent reduction in viability also in DLBCL cell lines, with no significant difference in sensitivity between ABC and GCB subtypes (Figure 1C). The calculated median IC50 values were 43.9 µM (95% CI: 10.7 - 77.1 µM) and 37.5 µM (95% CI: 15.5 - 59.6 µM) for ABC- and GCB-DLBCL cell lines. Overall, MZL and MCL cell lines exhibited greater sensitivity to CIGB-552 than DLBCL cell lines. When we compared cell lines with active (n = 9) or inactive p53 (n = 10), we only observed a trend toward greater CIGB-552 sensitivity in cell lines with functional p53 (Table S2, Figure S1).

To assess potential toxicity to non-malignant cells, we evaluated CIGB-552 activity in the non-malignant B- cell lymphoblastoid CB33 cell line using live-cell imaging. Cells were exposed to CIGB-552 at 6 µM for four days. No significant growth inhibition was observed in CB33 cells at the tested concentration (Figure S2).

Finally, given that canine lymphomas represent a relevant spontaneous model of the disease, we assessed the activity of CIGB-552 in the CLBL-1 canine DLBCL cell line. Using both Incucyte live-cell imaging and a 72-hour MTT assay. CIGB-552 showed dose-dependent cytotoxicity, with significant antiproliferative effects observed at concentrations as low as 0.3 µM in the Incucyte proliferation assay (Figure S13).

### CIGB-552 maintains dose-dependent antitumor activity in MZL cell lines with secondary resistance to BTK, PI3K, and BCL2 inhibitors

We next assessed anti-tumor activity in MZL cell lines with acquired secondary resistance to BTKi (VL51- Ibru (13)), BTKi/PI3Ki/BCL2i (Karpas1718-Ide (14)), PI3Ki (VL51-Ide (15), (SSK41-Copa)(17)), and PI3Ki/BCL2i (VL51-Copa (16), SSK41-CopaVene, SSK41-CopaVene14 (17)) and resistant clones with resistance to BCL2i (SSK41-Vene) (17). CIGB-552 maintained its dose-dependent effect across all cell lines, although activity was reduced in the resistant models derived from VL51 and Karpas1718 (Figure 1D, S3). Conversely, in SSK41, activity was higher in SSK41-CopaVene and SSK41-CopaVene14 than in their parental and other resistant cells (Figure 1D). Table 1 summarizes the IC50 values across all the B-cell lymphoma cell lines, showing the enrichment of MZL and MCL cells among the most sensitive cell lines, including the SSK41- resistant derivatives.

**Table 1.** Effect of CIGB-552 on the *in vitro* viability of B-cell lymphoma models. Cell lines were derived from marginal zone lymphoma (MZL), mantle cell lymphoma (MCL), and diffuse large B-cell lymphoma (DLBCL), and comprise drug-resistant derivatives. IC50 values were determined using the MTT assay after 72 h of exposure to CIGB-552. Results are based on at least two independent biological replicates.

| **Lymphoma cell lines** | **Histology** | **IC50 (µM)** |
|---|---|---|
| SSK41 C+V-Res | MZL | 12.77 |
| SSK41 CV14-Res | MZL | 13.01 |
| HC-1 | MZL | 15.16 |
| HAIRM | MZL | 16.74 |
| SP49 | MCL | 16.82 |
| SSK41 COP-Res | MZL | 17.24 |
| ESKOL | MZL | 18.18 |
| SSK41 | MZL | 18.24 |
| TMD8 | ABC-DLBCL | 22.36 |
| SSK41 VEN-Res | MZL | 23.30 |
| PFEIFFER | GCB-DLBCL | 24.39 |
| VL51 | MZL | 26.42 |
| Karpas1718 | MZL | 27.01 |
| CLBL1 | canine B cell lymphoma | 30.04 |
| OCI-LY7 | GCB-DLBCL | 30.79 |
| JEKO1 | MCL | 32.86 |
| MAVER1 | MCL | 33.90 |
| VL51 IBR-Res | MZL | 35.18 |
| Karpas1718 IDE-Res | MZL | 37.37 |
| VL51 COP-Res | MZL | 38.01 |
| REC1 | MCL | 43.38 |
| SU-DHL-2 | ABC-DLBCL | 43.93 |
| VAL | GCB-DLBCL | 44.32 |
| GRANTA-519 | MCL | 53.78 |
| VL51 IDE-Res | MZL | 59.84 |
| SU-DHL-16 | GCB-DLBCL | 74.96 |
| OCI-LY10 | ABC-DLBCL | 80.40 |

### CIGB-552 uptake varies across models and is associated with sensitivity

CIGB-552 enters cells through two mechanisms: energy-dependent endocytosis and direct membrane transduction (9). To distinguish between these pathways, we assessed peptide internalization at 37 °C (allowing both mechanisms) and 4 °C (avoiding endocytosis) in B cell lymphoma cell lines. Fluorescence- based internalization assays indicated that CIGB-552 uptake occurred predominantly through endocytosis in VL51 and SSK41 cells (both parental and drug-resistant derivatives) and JEKO-1 cells. In contrast, direct transduction appeared to contribute to peptide uptake in Karpas1718 and its derivative, as well as in HC- 1, ESKOL, HAIRM, and OCI-LY10 cells (Figure S4A-B).

Comparative analysis of CIGB-552 internalization across B-cell lymphoma cell lines revealed substantial variability (Figure 2A). SSK41-resistant derivatives showed significantly higher peptide internalization than parental cells. In contrast, internalization was higher in Karpas1718 than in its resistant derivative, whereas no significant difference was observed between parental VL51 cells and their ibrutinib-resistant derivative. Across the cell lines analyzed, CIGB-552 internalization negatively correlated with IC50 values (Pearson, *r* = −0.57, *p* = 0.02), indicating that higher peptide internalization was associated with greater sensitivity to CIGB-552 (Figure 2B).

**Figure 2.**
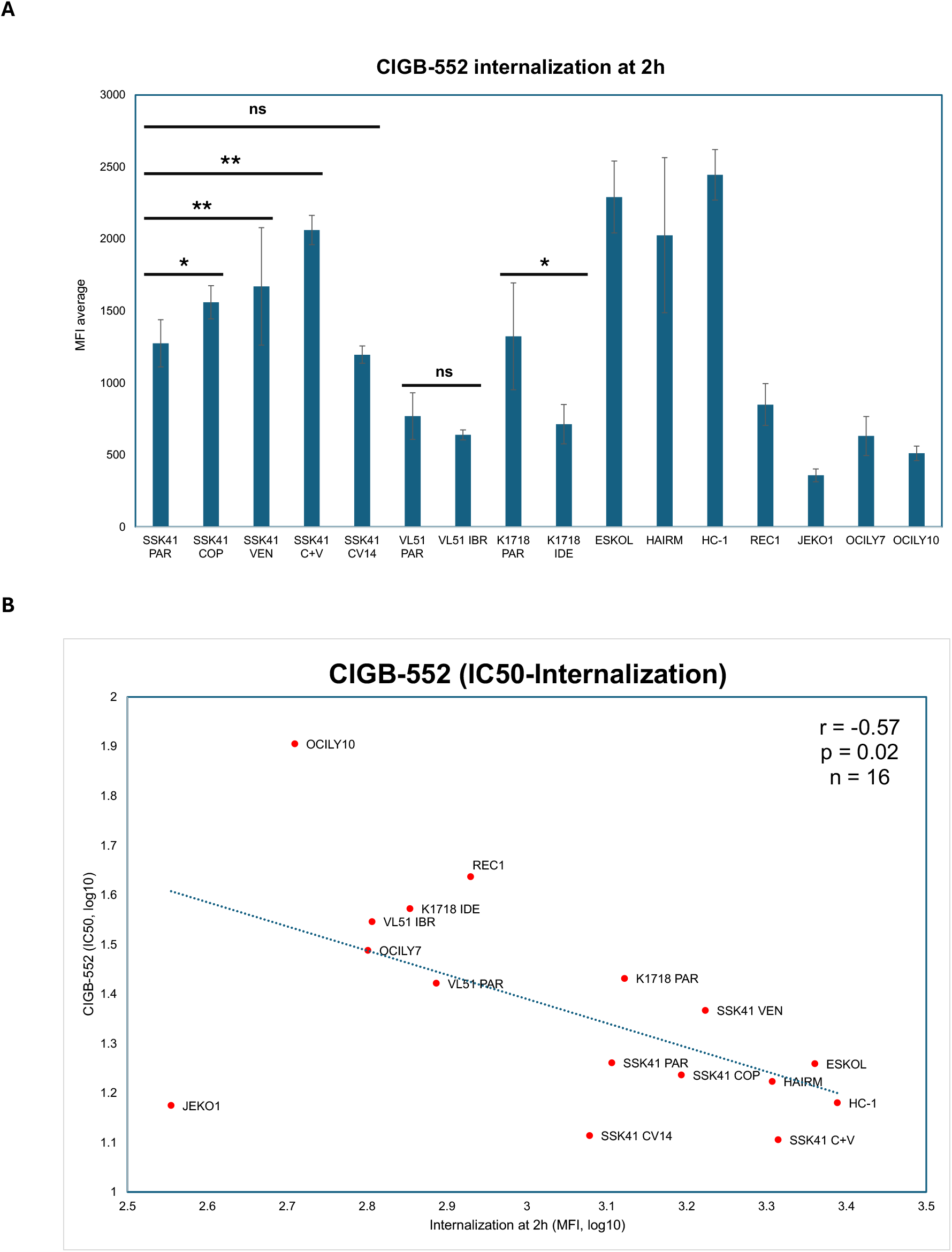
CIGB-552 internalization in B-cell lymphoma cell lines. (A) Parental MZL cell lines and their derivatives resistant to targeted agents were exposed to 5 μM fluorescein-labeled CIGB-552 for 2 h at 37°C. Peptide internalization was assessed by flow cytometry using fluorescein (FITC) and propidium iodide (PI) staining. PI-positive dead cells were excluded from the analysis. * indicates p values below 0.05, and ** indicates p values below 0.01 for comparisons of internalization between parental cell lines and their corresponding resistant clones. Bar plots represent the mean fluorescence intensity (MFI). Representative results from at least two independent experiments are shown. (B) CIGB-552 sensitivity is correlated with peptide internalization (Pearson’s r = −0.57, p = 0.02).

### CIGB-552 induces context-dependent cell death in B-cell lymphoma models

To characterize the mode of cell death caused by CIGB-552, apoptosis assays were conducted in MZL cell lines after 72 hours of treatment at twice the IC50. CIGB-552 induced apoptosis in VL51, ESKOL, and HAIRM, while necrosis was the dominant death mechanism in Karpas1718 and HC-1. Finally, in SSK41 cells, CIGB-552 induced both necrosis and apoptosis (Figure S5A).

Cell cycle analysis conducted after 24 hours of treatment with 50 µM CIGB-552 further supported these findings. An accumulation of cells in the Sub-G0 fraction was observed in VL51, ESKOL, HAIRM, and HC-1, whereas SSK41 cells showed G0/G1 arrest, and Karpas1718 cells accumulated in the G2/M phase (Figure S5B). This effect persisted when cells were exposed to twice the IC50 of CIGB-552 for 48 hours, with VL51 and Karpas1718 showing the same pattern, whereas SSK41 exhibited G2/M arrest (Figure S6).

To investigate the contribution of oxidative stress in the CIGB-552-induced cell death, Karpas1718 and SSK41 cells were pre-treated with vitamin E (α-tocopherol, 100 µM) prior to CIGB-552 exposure for 48 hours. Vitamin E partially rescued the anti-proliferative effect of CIGB-552 in Karpas1718 cells but had no protective effect in SSK41 cells (Figure S7), suggesting a cell line-specific reliance on oxidative stress mechanisms.

We also performed an apoptosis assay in MCL and DLBCL cell lines exposed to CIGB-552 or DMSO for 48 and 72 hours. In the MCL GRANTA-519 cell line, at 48 hours, the treated population showed a marked increase in both early (Annexin V+/PI-) and late (Annexin V+/PI+) apoptotic cells (Figure S8A), followed by a shift toward a higher necrotic population (PI+) and fewer cells in early/late apoptosis at 72 hours (Figure S8B). The GCB-DLBCL cell lines (PFEIFFER, SU-DHL-16, and OCI-LY7) exhibited predominant necrotic death. At 48 hours, a substantial proportion of cells were PI+ (necrotic), and at 72 hours, this population evolved toward an Annexin V+/PI+ profile, indicative of late-stage cell death (Figure S8A, B).

### CIGB-552 induces downregulation of oncogenic signaling and activation of stress response pathways

To gain mechanistic insight into the molecular effects of CIGB-552, we performed transcriptomic and proteomic profiling of MZL VL51 cells treated with CIGB-552 at the IC50 (26 µM) for 24 hours (Figure 3A-B, Table S3 and S4). Functional annotation of CIGB-552-modulated transcripts and proteins identified downregulation of key oncogenic pathways, including BCR signaling, NF-κB activation, PI3K/AKT/mTOR, TLR, MAPK, and JAK/STAT, as well as anti-apoptotic pathways (Figure 3C-D). Conversely, we observed enrichment of oxidative phosphorylation, fatty acid metabolism, DNA damage response, p53 signaling, protein ubiquitination, and stress responses. Taken together, these data provide a coherent molecular basis for the anti-lymphoma activity of CIGB-552, linking the suppression of pro-survival signals to metabolic reprogramming and DNA damage (Tables S5 and S6).

**Figure 3.**
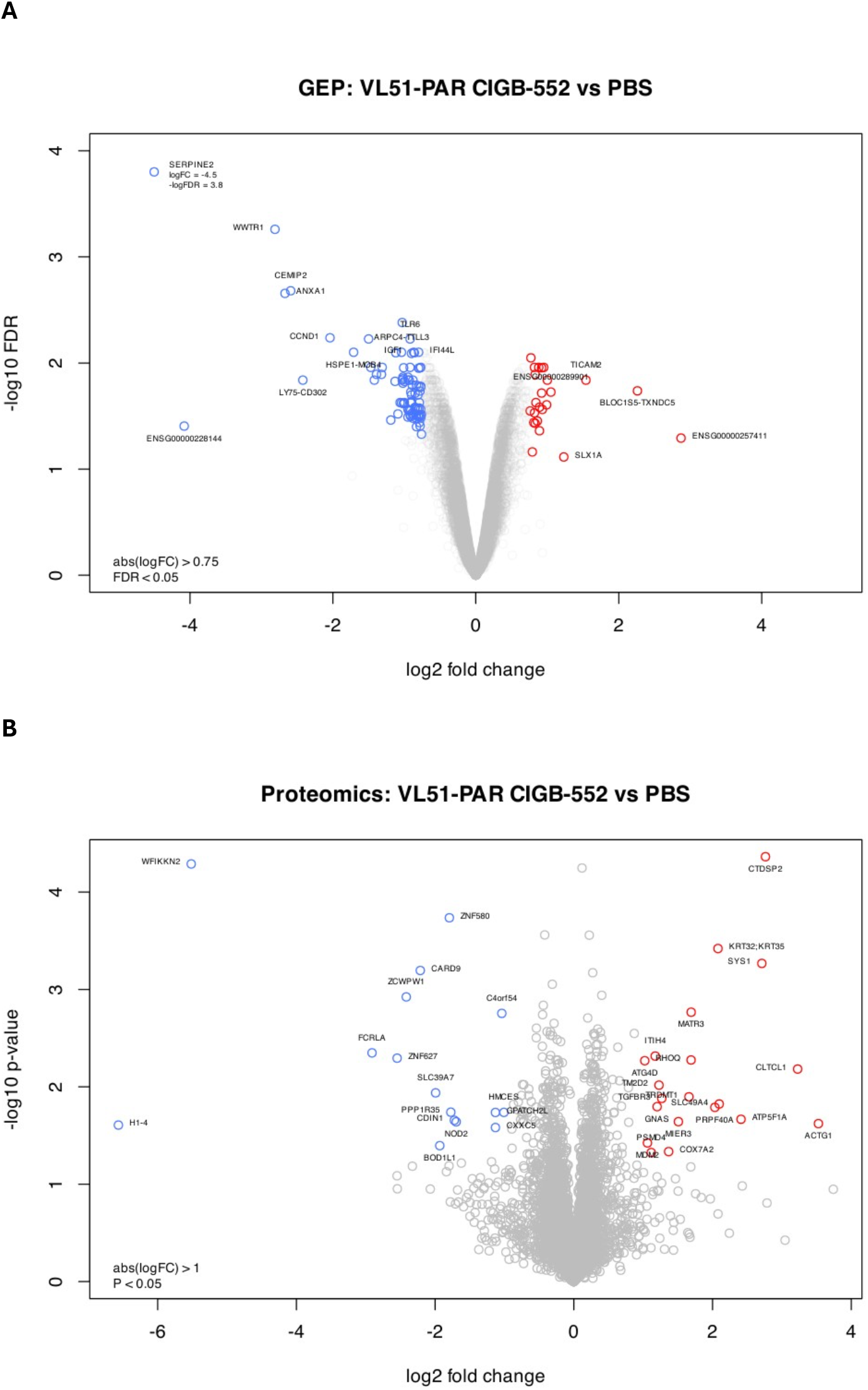
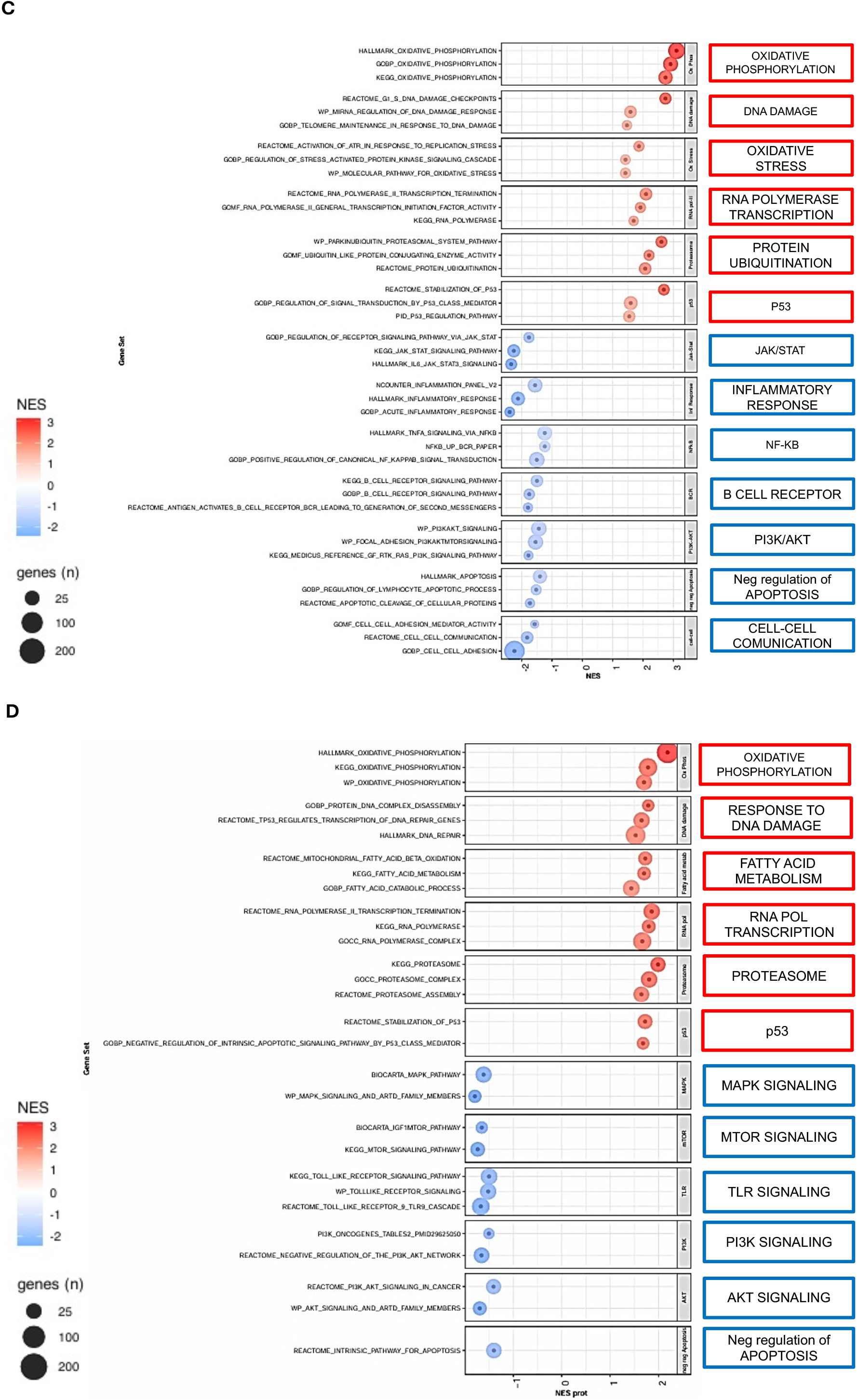
Transcriptomic and proteomic profiles of CIGB-552-treated VL51 cells. Cells were exposed to CIGB-552 for 24 h at the IC50 concentration (26 μM). Volcano plots show differential gene expression (RNA- seq, A) and differential protein abundance (mass spectrometry, B) following CIGB-552 treatment compared with PBS-treated controls. The x-axis represents the log₂ fold change (CIGB-552 vs. PBS). Differentially expressed genes and proteins are highlighted in red (upregulated) or blue (downregulated). Gene set enrichment analysis (GSEA) of the transcriptomic (C) and proteomic (D) profiles. Red dots indicate pathways with positive normalized enrichment scores (NES), reflecting enrichment in the CIGB-552-treated group, whereas blue dots indicate pathways with negative NES, reflecting enrichment in the PBS-treated control group.

### CIGB-552 modulates lymphoma survival proteins

Immunoblotting in VL51 cells confirmed the most relevant changes observed in the RNA and protein profiles, with reductions in NFκB1, NFκB2, AKT-S473 and ERK-Y204 phosphorylation, and cyclin D1 levels (Figure S9C). CIGB-552 reduced RelA levels as early as 4 hours post-treatment in SSK41 cells and decreased BCL2, MCL1, and AKT levels at 24 hours across MZL models (Figure S10A-C).

We also observed upregulation of COMMD1 protein levels after 24 hours of exposure to CIGB-552 in the VL51 cell line (Figure S9), in line with the reported mechanism of action of CIGB-552 (29).

PARP-1 protein levels showed a cell line-specific pattern that correlated with the mode of cell death: minimal change in VL51 (apoptosis), a decrease in SSK41 (apoptosis), and a marked increase in Karpas1718 (necrosis). This pattern was compatible with caspase-dependent cleavage in apoptotic cells (30) and PARP-1 hyperactivation in necrotic cell death (31,32), highlighting the context-dependent mechanism of CIGB-552.

### CIGB-552 enhances the antitumor activity of BTK inhibitors in parental and resistant MZL models

We next investigated whether the addition of CIGB-552 might enhance BTK inhibition, a clinically relevant therapeutic approach for patients with various B-cell lymphoid neoplasms, including MZL patients.

The combination of CIGB-552 with the BTK inhibitor ibrutinib was evaluated using both live-cell imaging (Incucyte) and metabolic read-out (MTT assay). VL51, K1718, and SSK41 cells and their respective derivatives with acquired resistance to BTK, PI3K, and BCL2 inhibitors were treated for 6 days with ibrutinib and CIGB-552. Combination treatment resulted in greater growth inhibition and lower cell viability than single agent across all cell lines (Figure 4, Figure S11). Synergy analysis using Chou–Talalay, Efficacy, HSA, and ZIP models consistently demonstrated a beneficial interaction between CIGB-552 and ibrutinib (Figure S11).

**Figure 4.**
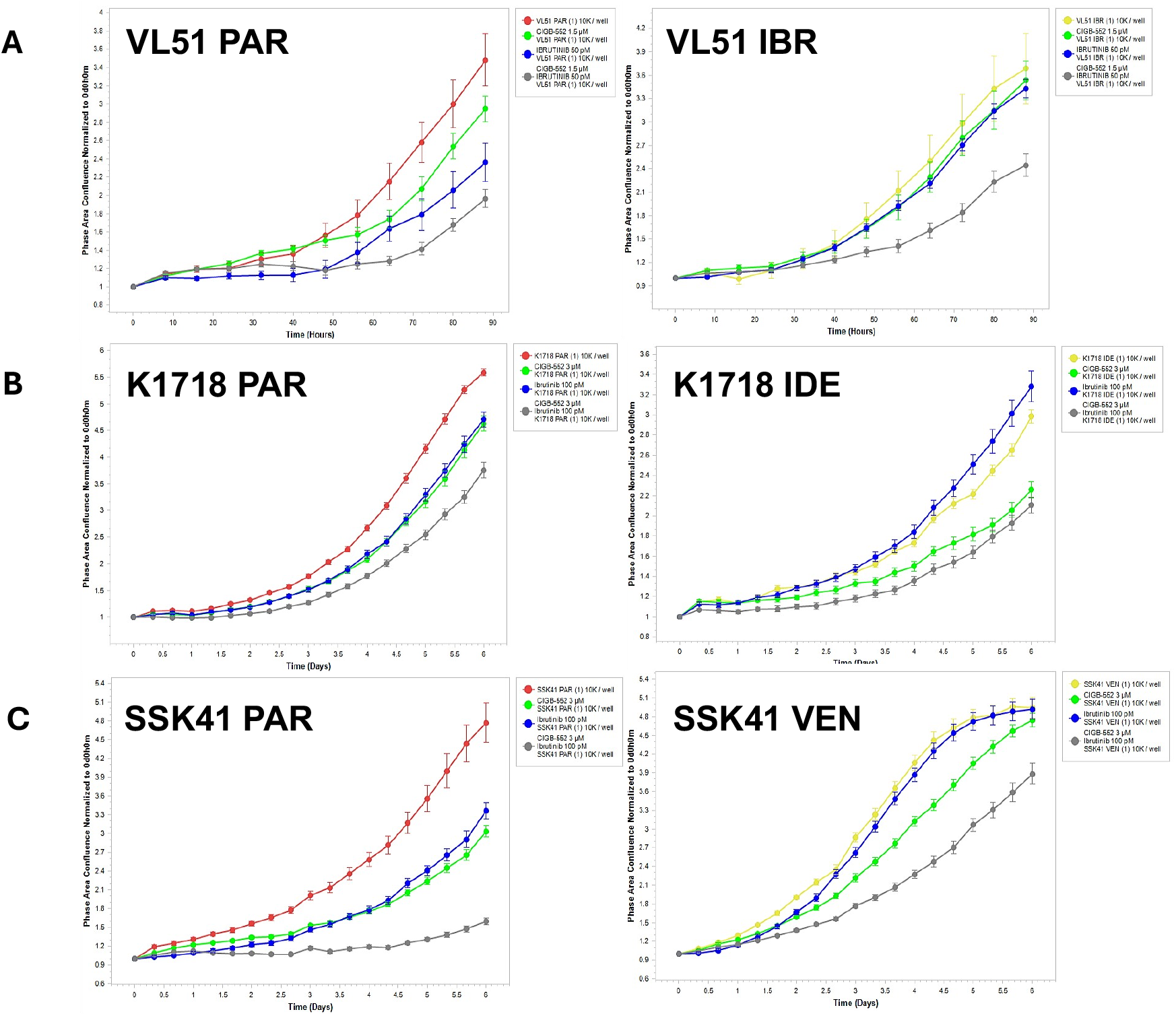
CIGB-552 enhances the effect of ibrutinib in MZL models with acquired resistance to PI3K, BTK, and BCL-2 inhibitors. The effect of the combination of ibrutinib and CIGB-552 on cell growth was assessed by real-time monitoring using the Incucyte system over 6 days of treatment. (A) Parental VL51 cells and their ibrutinib-resistant derivative were treated with ibrutinib (50 pM), CIGB-552 (1.5 μM), or the combination. (B, C) Parental and drug-resistant derivatives of Karpas1718 (B) and SSK41 (C) cells were treated with ibrutinib (100 pM), CIGB-552 (3 μM), or the combination. Representative results from two independent biological replicates are shown.

Furthermore, these findings were extended by combining CIGB-552 with other BTK-targeting agents in the SSK41 cell line: the second-generation covalent inhibitor zanubrutinib, the non-covalent inhibitor pirtobrutinib, and the BTK degrader bexobrutideg (NX-5948). The addition of the peptide enhanced the cytotoxicity of all compounds (Figure S12), supporting the broad combinability of CIGB-552 with BTK- targeting therapies.

Finally, we also validated the benefit of combining CIGB-552 with ibrutinib in the canine model CLBL-1 (Figure S13A-B) using Incucyte and MTT assays. In both assays, the combination treatment resulted in significantly greater growth inhibition than either agent alone.

### CIGB-552 exhibits cytotoxic activity in primary B cells and enhances the activity of ibrutinib in primary MZL cells

The cytotoxic activity of CIGB-552 was evaluated by Incucyte in primary cells from patients with MZL, MCL, relapsed/refractory MCL, and DLBCL (Figure 5A). CIGB-552 treatment resulted in reduced cell growth compared with DMSO-treated controls across the primary lymphoma samples (Figure 5B), supporting the activity of CIGB-552 in patient-derived cells. Finally, we evaluated whether CIGB-552 could enhance the activity of BTK inhibition in primary MZL cells. The combination of CIGB-552 and ibrutinib resulted in greater growth inhibition than either single treatment. These results further support our previous *in vitro* findings that CIGB-552 enhances ibrutinib activity in MZL cells (Figure 5C).

**Figure 5.**
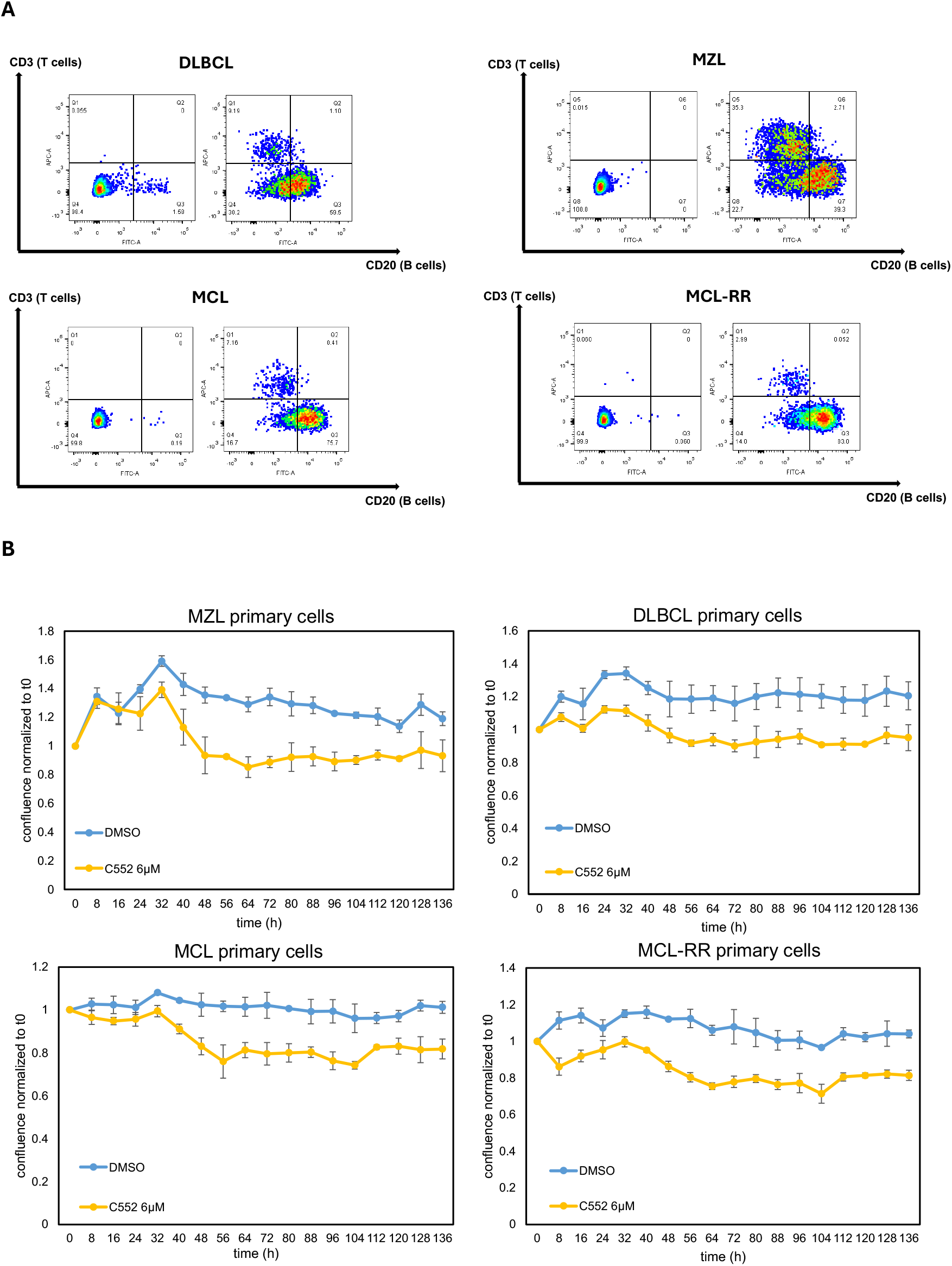
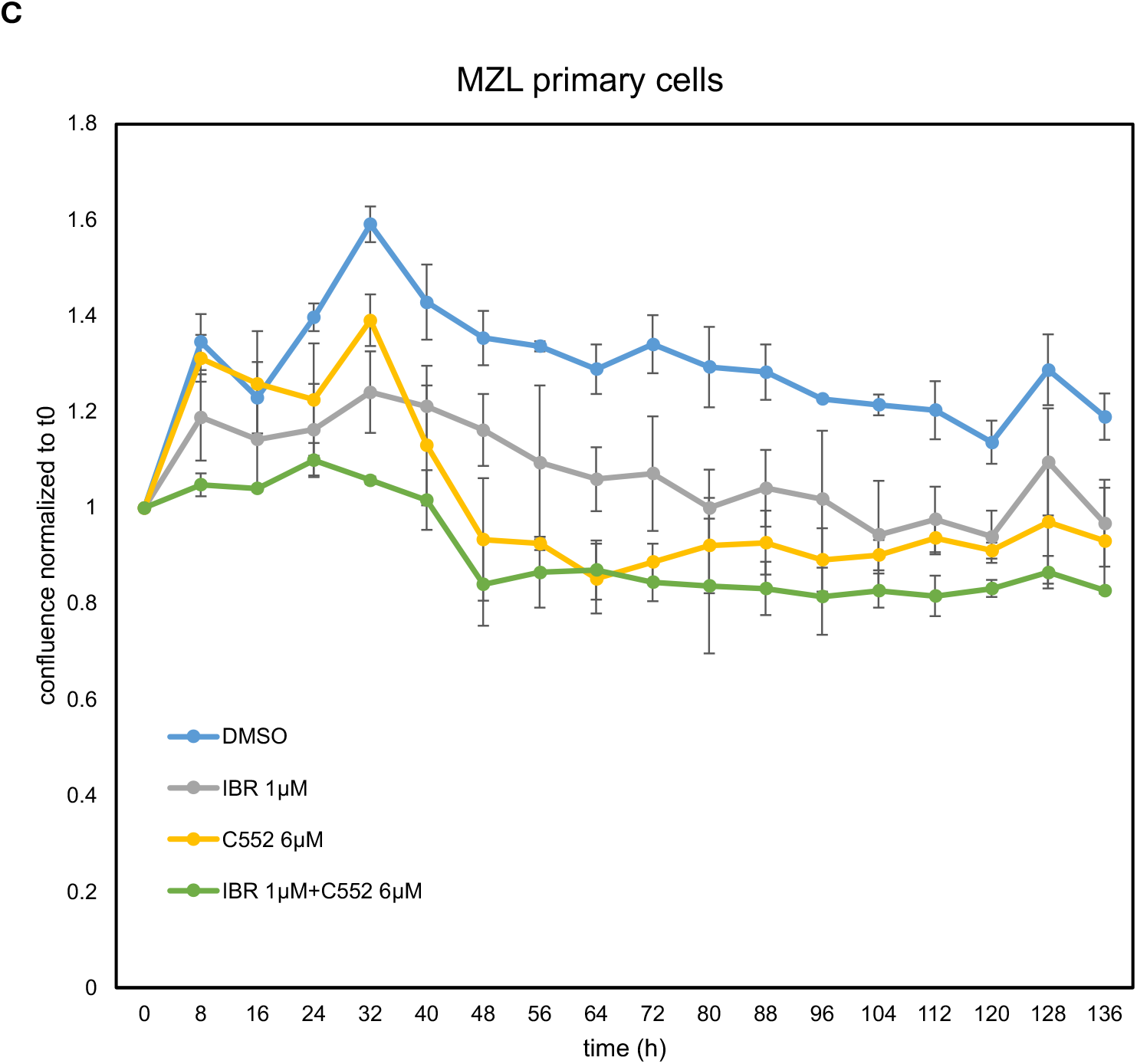
Antitumor activity of CIGB-552 in primary lymphoma cells. Primary cells from patients with marginal zone lymphoma (MZL), mantle cell lymphoma (MCL), relapsed/refractory MCL (MCL-RR), and diffuse large B-cell lymphoma (DLBCL) were treated with CIGB-552 as a single agent, and cell growth was monitored by live-cell imaging using the Incucyte system. (A) Flow cytometry analysis of CD20 and CD3 staining showed that most cells in the primary samples were B cells. (B) Primary lymphoma cells were treated with CIGB-552 (6 μM) as a single agent. (C) Primary MZL cells were treated with CIGB-552 (6 μM) in combination with ibrutinib (1 μM). Cell growth was monitored using the Incucyte system.

## Discussion

This study extends the preclinical evaluation of CIGB-552 to B-cell lymphomas. The peptide reduced viability in MZL, MCL, and DLBCL cell lines and retained activity in MZL derivatives with acquired resistance to several targeted agents. Importantly, CIGB-552 enhanced the effects of BTK-directed therapies, and activity was also observed in primary lymphoma samples. Together, these findings provide a rationale for further evaluation of CIGB-552 in B-cell malignancies.

The response of drug-resistant models is particularly relevant. Although CIGB-552 remained active across the resistant derivatives tested, some VL51- and Karpas1718-derived models were less sensitive than their parental counterparts, whereas selected SSK41-resistant derivatives were more sensitive. Thus, the data indicate retained, albeit heterogeneous, activity reversal of resistance. Determining which resistance- associated adaptations can be overcome by the peptide will be important for future studies.

Activity was observed across lymphoma histologies, although MZL and MCL models generally showed lower IC50 values than DLBCL models. Even though the modest number of cell lines within each subgroup and the variability between models preclude firm conclusions about histology-specific sensitivity, the distribution of responses supports prioritizing MZL and MCL for further investigation.

CIGB-552 uptake differed across cell lines and was associated with drug sensitivity, consistent with cellular delivery contributing to response. Temperature-dependent internalization assays suggested an important role for endocytosis in several models but did not establish a single uptake mechanism across all lymphoma cells. Future studies should determine whether uptake predicts response in primary specimens and whether specific transport pathways can be manipulated.

CIGB-552 induced distinct cell-death phenotypes, including apoptotic and non-apoptotic responses, and altered cell-cycle distributions. Partial protection by the antioxidant vitamin E (α-tocopherol) in selected models supports a contribution of oxidative stress, although the present experiments do not define the complete sequence of events leading to cell death. These differences suggest that cellular context influences the response to the peptide.

Transcriptomic and proteomic profiling, and immunoblotting analyses indicated broad changes in pathways relevant to lymphoma survival, including BCR, NF-κB, PI3K/AKT/mTOR, MAPK, and JAK/STAT signaling, and reduced abundance or activation of key effectors, including AKT, ERK, cyclin D1, BCL2, and MCL1. Because the omics experiments were concentrated in one cell line, these findings should be interpreted as treatment-associated signatures. CIGB-552 also increased COMMD1 protein abundance in VL51 cells, consistent with observations in other tumor models (33–36).

The downregulation of cyclin D1 is also consistent with the strong effect observed in MCL cell lines, which harbor the t(11;14) translocation leading to constitutive cyclin D1 overexpression. Interestingly, FCRLA, a protein implicated in B-cell differentiation and lymphoma biology, was downregulated at both the protein and transcript levels, suggesting a potential role for FCRLA in the response to CIGB-552. SERPINE2 was the most strongly downregulated gene following CIGB-552 treatment, highlighting it as a potential candidate for further investigation in B-cell lymphoma.

The combination experiments provide a clinically relevant direction for further development. CIGB-552 increased the effects of ibrutinib in parental and resistant MZL models and showed favorable interactions with zanubrutinib, pirtobrutinib, and a BTK degrader in the models tested. These results suggest complementarity between peptide-induced cellular perturbations and BTK-directed treatment. Indeed, CIGB-552-suppressed pathways that overlap with known adaptive circuits sustaining survival despite BTK inhibition (37), particularly NF-κB, PI3K/AKT, and MAPK signaling.

The observation of activity in primary lymphoma samples strengthens the translational relevance of the cell-line results. In primary MZL samples, the combination with ibrutinib further reduced growth compared with the individual agents. The number of specimens remains limited, however, and additional samples, ideally including treatment-exposed disease, will be required to establish the consistency and magnitude of these effects.

The canine CLBL-1 model provided an additional comparative system in which CIGB-552 inhibited growth and enhanced ibrutinib activity. These results broaden the experimental observations but should not be taken as a substitute for *in vivo* efficacy and tolerability studies. Given the similarities between the clinical features of canine and human lymphomas (38–40), including the early development of BTK inhibitors in dogs with spontaneous lymphomas (41), this result provides additional support for the clinical exploration of CIGB-552, both as a single agent and in rational combination strategies, in both the human and the canine contexts.

In summary, CIGB-552 shows preclinical activity in B-cell lymphomas, including models with acquired resistance to targeted agents, and enhances the effects of BTK-directed therapies. Further work should define the determinants of peptide uptake and response, establish the contribution of COMMD1, and evaluate efficacy and safety in relevant *in vivo* models before clinical translation.

## Authorship

M.C.A. and A.J.A. designed and performed experiments, interpreted data, and co-wrote the manuscript; A.A.M designed experiments and interpreted data; E.P. performed experiments; F.S. provided advice, and performed experiments; A.R. performed transcriptome profiling; M.P. performed proteomic profiling; A.J.A. and L.C. performed bioinformatics analyses; L.A., provided advice and resources; R.M., N.M. and G.G.: collected and annotated clinical specimens; F.Si., provided resources; F.C., provided advice; J.R.F. co- designed the study and provided compounds; F.B. co-designed the study, interpreted data, and co-wrote the manuscript. All Authors have approved the final manuscript.

