## Supplementary tables and figures for "The antitumor peptide CIGB-552 is active in B-cell lymphomas and enhances BTK-targeted therapies"

**<u>Running title</u>**

CIGB-552 in B-cell lymphomas

**Supplementary tables**

**Table S1.** Panel of proteins tested in the immunoblotting experiments

| **source** | **protein** | **company** | **# catalog** |
|---|---|---|---|
| rabbit polyclonal | α-AKT | Cell Signaling | 9272 |
| rabbit polyclonal | α-p(S473) AKT | Cell Signaling | 4060 |
| mouse monoclonal | α-vinculin | Santa Cruz Biotechnology | 25336 |
| rabbit monoclonal | RELA NF-kB/p65 (C22B4) | Cell Signaling | 4764 |
| rabbit monoclonal | p(S536)-NF-kappaB p65 | Cell signaling | 3033 |
| rabbit monoclonal | NF-kB1 p105/p50 (D4P4D) | Cell Signaling | 13586 |
| rabbit polyclonal | NF-kB2 p100/p52 | Cell Signaling | 4882 |
| mouse polyclonal | COMMD1 | SIGMA | 1408701 |
| mouse monoclonal | Cyclin D1 (DCS-6) | Santa Cruz Biotechnology | 20044 |
| mouse monoclonal | ERK1 (G-8) | Santa Cruz Biotechnology | 271269 |
| mouse monoclonal | p(Y204) ERK1 | Santa Cruz Biotechnology | 7383 |
| mouse monoclonal | PARP-1 (F-2) | Santa Cruz Biotechnology | 8007 |
| mouse monoclonal | Bcl-2 (C-2) | Santa Cruz Biotechnology | 7382 |
| rabbit monoclonal | MCL-1 (D35A5) | Cell signaling | 5453S |

**Table S2. Lymphoma cell lines histology, cell of origin, doubling time (hours) and genomic annotation.** DLBCL, diffuse large B-cell lymphoma; GCB, germinal center B-cell; ABC, activated B-cell; MCL, mantle cell lymphoma; MZL, marginal zone lymphoma; cDLBCL, canine diffuse large B-cell lymphoma. Data from (1) and https://www.cellosaurus.org/CVCL_L322.

| Cell line | Histology | Cell of origin | Doubling time (hours) | p53 |
|---|---|---|---|---|
| SU-DHL-16 | DLBCL | GCB | 40 | active |
| VAL | DLBCL | GCB | 60 | active |
| Pfeiffer | DLBCL | GCB | 24-30 | inactive |
| OCI-Ly7 | DLBCL | GCB | 19 | inactive |
| SU-DHL-2 | DLBCL | ABC | 24 | active |
| OCI-Ly10 | DLBCL | ABC | 95 | inactive |
| TMD8 | DLBCL | ABC | 24-30 | active |
| GRANTA-519 | MCL | - | 48 | inactive |
| JeKo-1 | MCL | - | 26-50 | inactive |
| MAVER-1 | MCL | - | 24-48 | inactive |
| REC-1 | MCL | - | 38 | inactive |
| SP-49 | MCL | - | 60 | active |
| KARPAS-1718 | MZL | - | 35-50 | inactive |
| SSK41 | MZL | - | 30-45 | inactive |
| VL51 | MZL | - | 70-80 | active |
| ESKOL | MZL | - | NA | active |
| HAIR-M | MZL | - | 48 | active |
| HC-1 | MZL | - | 45 | active |
| CLBL1 | cDLBCL | - | 19 | inactive |

**Table S3. Differential gene expression analysis of CIGB-552-treated VL51 cells using limma.** Results of differential gene expression analysis comparing VL51 cells treated with CIGB-552 (26 μM, 24 h) with untreated control cells. Statistical analysis was performed using the limma package. RNA-seq data are publicly available in the Gene Expression Omnibus (GEO) repository under accession number GSE354291.

**Table S4. Differential protein expression analysis of CIGB-552-treated VL51 cells using limma.** Results of differential protein abundance analysis comparing VL51 cells treated with CIGB-552 (26 μM, 24 h) with untreated control cells. Statistical analysis was performed using the limma package.

**Table S5.** Top 10 upregulated and downregulated genes identified by RNA-seq following treatment of the VL51 MZL cell line with CIGB-552 at its IC50 for 24 h. Statistically significant *p*-values and adjusted *p*-values (*q*-values) ($p < 0.05$ and $q < 0.05$) are highlighted in red. Gene descriptions were obtained from GeneCards and UniProtKB.

| gene name | protein name | logFC | P.Value | adj.P.Val | description |
|---|---|---|---|---|---|
| **TICAM2** | TIR domain-containing adapter molecule 2 | 1.54 | <0.01 | 0.01 | Sorting adaptor protein involved in Toll-like receptor signaling and downstream innate immune responses, including type I interferon induction. |
| **PCTP** | Phosphatidylcholine transfer protein | 1.00 | <0.01 | 0.01 | Lipid-transfer protein that binds phosphatidylcholine and mediates its transfer between membranes, contributing to phospholipid transport. |
| **GCSH** | Glycine cleavage system H protein, mitochondrial | 0.92 | <0.01 | 0.01 | Mitochondrial component of the glycine cleavage system that transfers the methylamine group during glycine degradation. |
| **RPP40** | Ribonuclease P protein subunit p40 | 0.89 | <0.01 | 0.03 | Core subunit of the RNase P and RNase MRP complexes, involved in tRNA 5'-leader removal and precursor RNA processing. |
| **LDHC** | L-lactate dehydrogenase C chain | 0.89 | <0.01 | 0.04 | NAD-dependent lactate dehydrogenase isoenzyme associated with the reversible conversion of lactate and pyruvate; predominantly expressed in the male reproductive system. |
| **MOSPD1** | Motile sperm domain-containing protein 1 | 0.86 | <0.01 | 0.03 | Predicted membrane-associated protein implicated in cell differentiation and the regulation of RNA polymerase II-mediated transcription. |
| **PDCL3** | Phosducin-like protein 3 | 0.83 | <0.01 | 0.01 | Phosducin-like protein with chaperone functions, including regulation of VEGFR2 abundance, protein folding, cytoskeletal actin folding, and caspase activation during apoptosis. |
| **MATR3** | Matrin-3 | 0.82 | <0.01 | 0.01 | Nuclear matrix protein involved in transcriptional regulation and RNA processing, retention, and quality control; also contributes to the stability of m6A-containing MYC mRNA. |
| **LYRM4** | LYR motif-containing protein 4 | 0.76 | 0.001 | 0.03 | Mitochondrial iron–sulfur cluster assembly factor that stabilizes the core assembly complex and supports the formation of iron–sulfur clusters. |
| **EAF2** | ELL-associated factor 2 | 0.75 | 0.002 | 0.03 | Transcription elongation factor that promotes RNA polymerase II-mediated transcription through interactions with ELL-family elongation factors. |
| **IL13RA1** | Interleukin-13 receptor subunit alpha-1 | -1.02 | <0.01 | 0.01 | Cell-surface cytokine receptor subunit that binds IL-13 and participates in IL-13 and IL-4 receptor signaling. |

| | | | | | |
|---|---|---|---|---|---|
| **CDC42BPA** | Serine/threonine-protein kinase MRCK alpha | -1.02 | <0.01 | 0.01 | Serine/threonine kinase acting downstream of CDC42 to regulate actin cytoskeleton organization, cell migration, and cellular extrusion. |
| **ST3GAL1** | CMP-N-acetylneuraminate-beta-galactosamide-alpha-2,3-sialyltransferase 1 | -1.06 | <0.01 | 0.02 | Glycosyltransferase that catalyzes α2,3-sialylation of glycoproteins and glycolipids, contributing to cell-surface glycan modification. |
| **PMEPA1** | Protein TMEPAI | -1.39 | <0.01 | 0.01 | Transmembrane adaptor protein involved in negative feedback regulation of TGF-β and androgen receptor signaling, including proteasome-mediated degradation of the androgen receptor. |
| **ITM2B** | Integral membrane protein 2B | -1.47 | <0.01 | 0.01 | Integral membrane protein that regulates amyloid precursor protein processing and inhibits amyloid-beta aggregation; also associated with neurite outgrowth. |
| **CCND1** | G1/S-specific cyclin-D1 | -2.04 | <0.01 | <0.01 | Cyclin that forms regulatory complexes with CDK4/CDK6 to promote cell-cycle progression through the G1/S transition. |
| **CEMIP2** | Cell surface hyaluronidase CEMIP2 | -2.59 | <0.01 | <0.01 | Cell-surface hyaluronidase that cleaves extracellular hyaluronan into intermediate-sized fragments; involved in extracellular matrix remodeling and regulation of VEGF-associated angiogenesis. |
| **ANXA1** | annexin A1 | -2.67 | <0.01 | <0.01 | Calcium-dependent phospholipid-binding protein involved in the regulation of inflammation, glucocorticoid-mediated responses, phagocytosis, and antitumor immune responses. |
| **WWTR1** | WW domain-containing transcription regulator protein 1 | -2.81 | <0.01 | <0.01 | Transcriptional coactivator and downstream effector of the Hippo signaling pathway, involved in the regulation of cell proliferation, apoptosis, and differentiation. |
| **SERPINE2** | serpin family E member 2 | -5.73 | <0.01 | <0.01 | Serine protease inhibitor involved in the regulation of extracellular proteolysis; also binds heparin and can promote neurite extension. |

**Table S6.** Top 10 upregulated and downregulated proteins identified by Proteomic following treatment of the VL51 MZL cell line with CIGB-552 at its IC50 for 24 h. Statistically significant *p*-values and adjusted *p*-values (*q*-values) ($p < 0.05$ and $q < 0.05$) are highlighted in red. Gene descriptions were obtained from GeneCards and UniProtKB.

| gene name | protein name | logFC | P.value | adj.P.Val | decription |
|---|---|---|---|---|---|
| **ACTG1** | Actin, cytoplasmic 2 | 3.53 | 0.02 | 0.28 | Cytoplasmic actin involved in cytoskeletal organization and intracellular cell motility. |
| **CLTCL1** | Clathrin heavy chain 2 | 3.23 | 0.01 | 0.01 | Clathrin-related protein involved in coated-vesicle formation and intracellular trafficking. |
| **CTDSP2** | Carboxy-terminal domain RNA polymerase II polypeptide A small phosphatase 2 | 2.76 | <0.01 | 0.00 | Phosphatase that regulates RNA polymerase II transcription by dephosphorylating its C-terminal domain. |
| **SYS1** | Protein SYS1 homolog | 2.71 | <0.01 | 0.00 | Golgi-associated protein involved in intracellular protein trafficking. |
| **ATP5F1A** | ATP synthase F(1) complex subunit alpha, mitochondrial | 2.41 | 0.02 | 0.29 | Mitochondrial ATP synthase subunit involved in ATP production through oxidative phosphorylation. |
| **SLC49A4** | Solute carrier family 49 member 4 | 2.10 | 0.01 | 0.27 | Membrane transporter involved in proton-dependent pyridoxine (vitamin B6) transport. |
| **PRPF40A** | Pre-mRNA-processing factor 40 homolog A | 2.03 | 0.02 | 0.27 | Nuclear scaffolding protein involved in pre-mRNA splicing and regulation of cytoskeletal dynamics. |
| **MATR3** | Matrin-3 | 1.69 | <0.01 | 0.01 | Nuclear matrix protein involved in cell morphology, cytoskeletal organization, and RNA processing. |
| **RHOQ** | Rho-related GTP-binding protein RhoQ | 1.69 | 0.01 | 0.08 | Small GTPase involved in intracellular signal transduction and epithelial cell polarization. |
| **TRDMT1** | tRNA (cytosine(38)-C(5))-methyltransferase | 1.66 | 0.01 | 0.26 | RNA methyltransferase that specifically methylates cytosine 38 in the anticodon loop of tRNA-Asp. |
| **PPP1R35** | Protein phosphatase 1 regulatory subunit 35 | -1.77 | 0.02 | 0.30 | Regulatory protein involved in centriole elongation and centrosome biogenesis. |
| **ZNF580** | Zinc finger protein 580 | -1.79 | <0.01 | <0.01 | Zinc finger transcription factor involved in transcriptional regulation and associated with endothelial cell proliferation and inflammatory responses. |
| **BOD1L1** | Biorientation of chromosomes in cell division protein 1-like 1 | -1.93 | 0.04 | 0.49 | Chromosome-associated protein involved in DNA damage response, replication fork protection, and DNA repair. |
| **SLC39A7** | Zinc transporter SLC39A7 | -1.99 | 0.01 | 0.19 | Zinc transporter that mobilizes $Zn^{2+}$ from the endoplasmic reticulum and Golgi apparatus to the cytosol, contributing to intracellular zinc homeostasis. |
| **CARD9** | Caspase recruitment domain-containing protein 9 | -2.21 | 0.00 | 0.00 | Signaling adaptor that promotes NF-κB and MAPK activation downstream of innate immune receptors. |
| **ZCWPW1** | Zinc finger CW-type PWWP domain protein 1 | -2.42 | 0.00 | 0.00 | Histone methylation reader involved in chromatin regulation and PRDM9-associated meiotic DNA double-strand break repair. |
| **ZNF627** | Zinc finger protein 627 | -2.55 | 0.01 | 0.01 | KRAB domain-containing zinc finger transcription factor predicted to regulate RNA polymerase II-dependent transcription. |
| **FCRLA** | Fc receptor-like A | -2.91 | <0.01 | 0.01 | B-cell-associated immunoglobulin-like protein implicated in B-cell differentiation and lymphoma biology. |
| **WFIKKN2** | WAP, Kazal, immunoglobulin, Kunitz and NTR domain-containing protein 2 | -5.52 | <0.01 | <0.01 | Extracellular protease inhibitor that also modulates myostatin activity and TGF-β-related signaling. |
| **H1-4** | Histone H1.4 | -6.57 | 0.02 | 0.26 | Linker histone involved in chromatin organization, nucleosome spacing, and transcriptional regulation. |

### Supplementary Figures

**Supplementary Figure S1. Association between p53 functional status and CIGB-552 sensitivity in parental B-cell lymphoma cell lines.** Dot plot showing the distribution of CIGB-552 $IC_{50}$ values in parental B-cell lymphoma cell lines (MZL, MCL, and DLBCL) grouped according to p53 functional status (active or inactive). Each dot represents an individual cell line, and horizontal lines indicate the median $IC_{50}$ values. Although cell lines with functional p53 tended to exhibit lower $IC_{50}$ values, the difference between groups was not statistically significant (two-tailed unpaired t-test with Welch's correction, $p = 0.4696$).

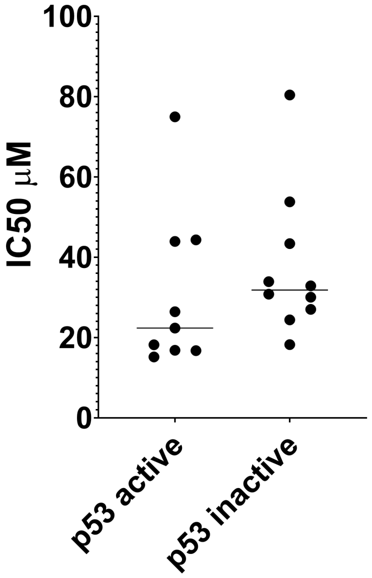

**Supplementary Figure S2. Antitumor activity of CIGB-552 in non-malignant B cell lymphoblastoid cell line**. Pharmacological profile of CIGB-552 at 6 μM on CB33 non-malignant B cell lymphoma cell line, evaluated by Incucyte (4 days of exposure), DMSO was used as control.

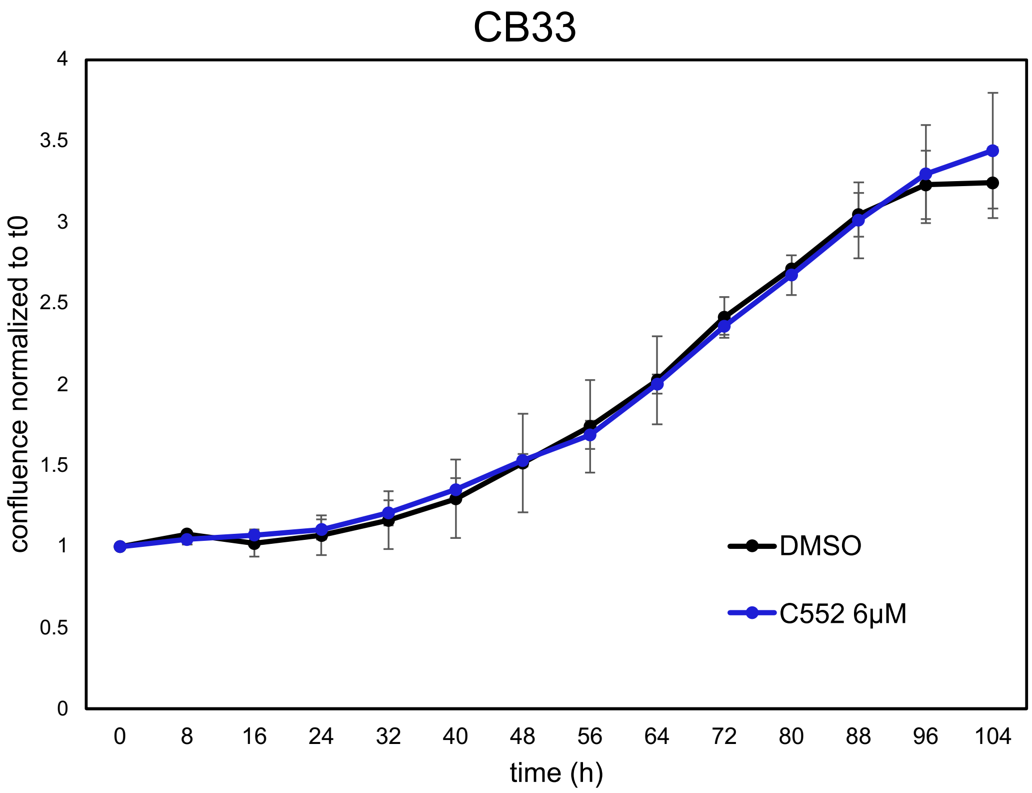

**Supplementary Figure S3. Pharmacological profile of CIGB-552 in MZL parental and resistant cell lines.** K1718 (PAR and IDE-Res), VL51 (PAR and IBR-Res), and SSK41 (PAR and VEN-Res) cell lines were evaluated by Incucyte (6 days of exposure) upon the treatment with CIGB-552 at different concentrations. DMSO was used as control.

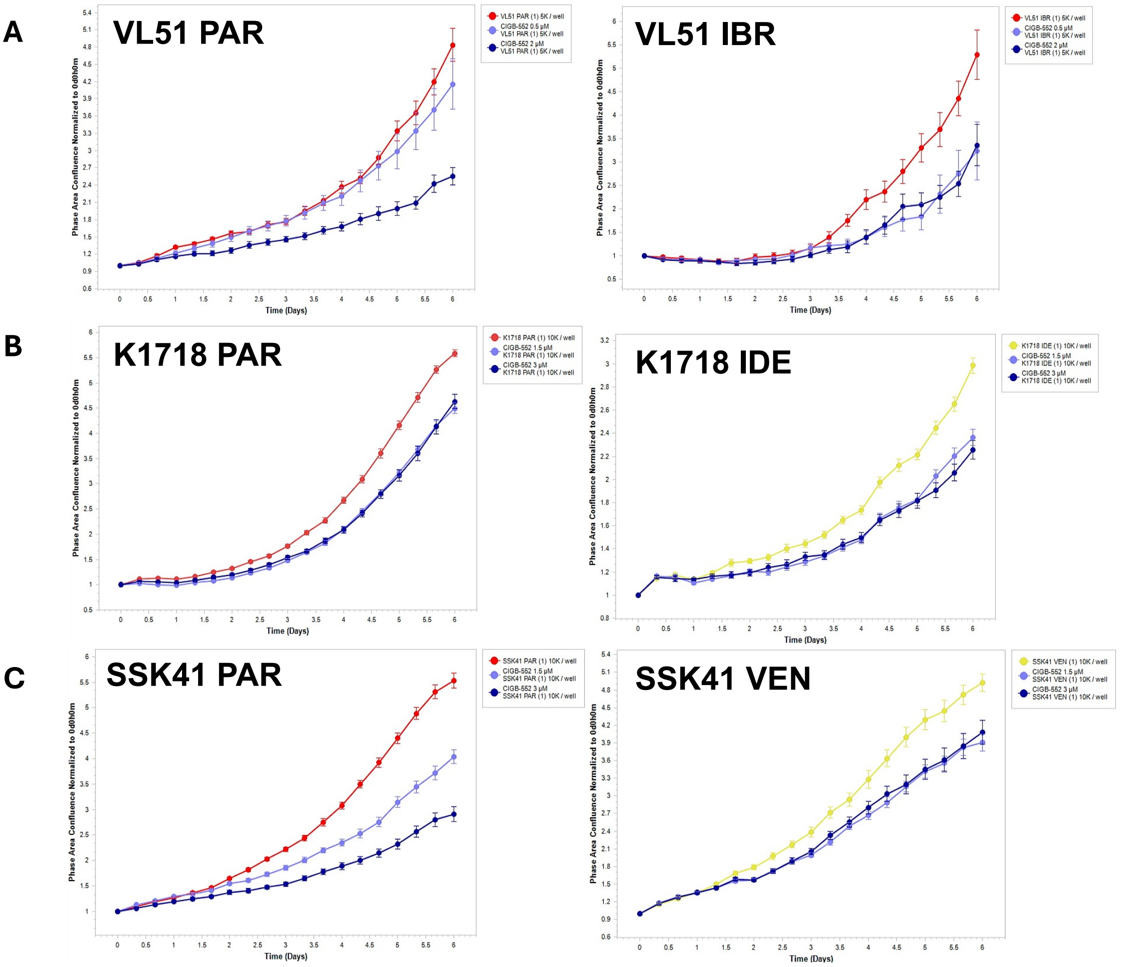

**Supplementary Figure S4. CIGB-552 internalization in B-cell lymphoma cell lines and the effect of temperature.** (A) Cells were exposed to 5 μM fluorescein-labeled CIGB-552 for 1 h at 37°C (orange curves) or 4°C (blue curves). Peptide-associated fluorescence was assessed by flow cytometry using FITC and propidium iodide (PI) staining. (B) Heatmap summarizing CIGB-552 internalization, the estimated relative contributions of endocytosis and direct transduction, COMMD1 mRNA expression (RNA-seq), and sensitivity to CIGB-552 (IC50) across parental and drug-resistant B-cell lymphoma cell lines. Colors indicate relative values within each column, with red representing higher and blue representing lower values. SSK41 VEN-Res cells showed higher CIGB-552 internalization than parental SSK41 cells ($p$ = 0.04). Representative results from at least two independent experiments are shown.

**A**

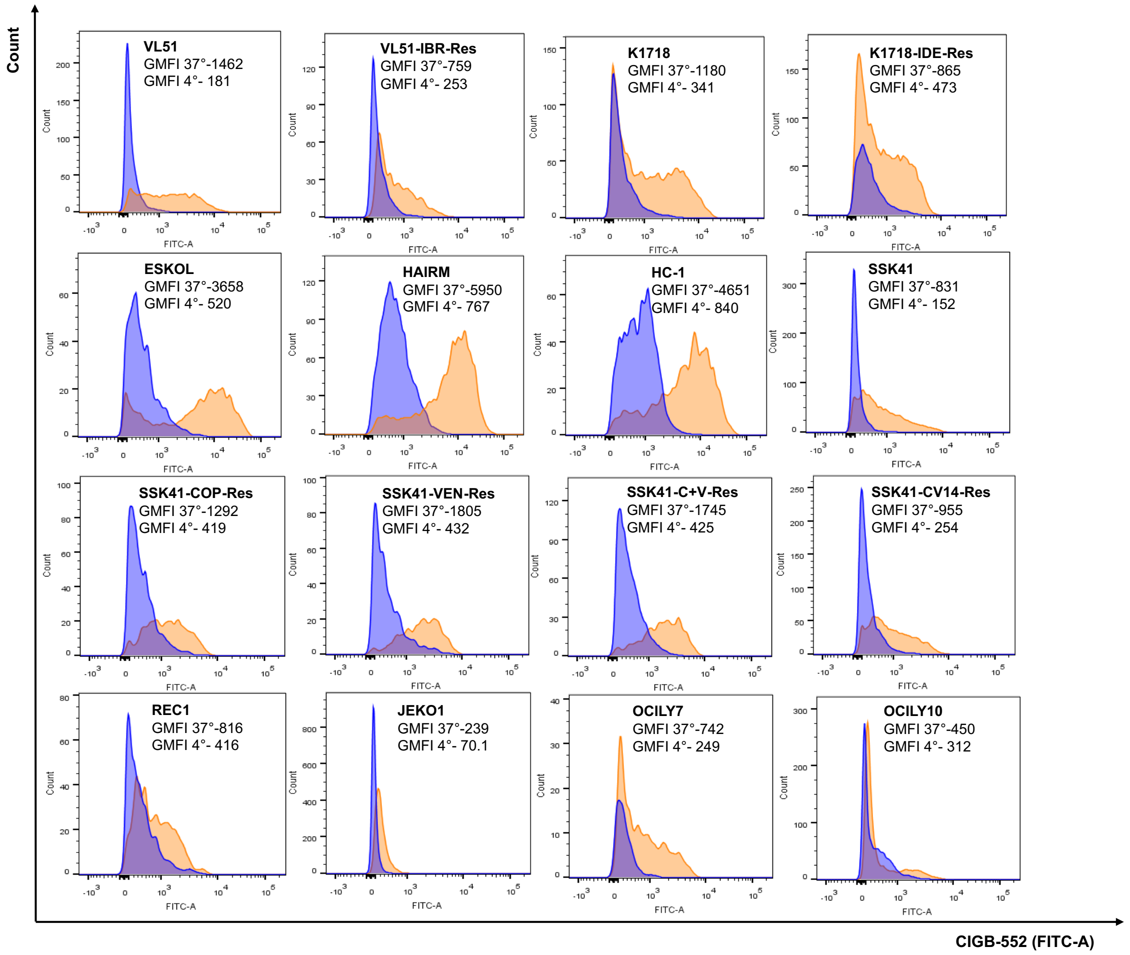

**B**

| Cell lines | Type | IC50 (μM ) | Internalization at 2h GMFI | % of transduction | % of endocytosis | Commd1 (lcpm, RNA seq) |
|---|---|---|---|---|---|---|
| **SSK41-CopaVene** | MZL | 12.77 | 2062.26 | 34.02 | 65.98 | 5.14 |
| **SSK41-CopaVene14** | MZL | 13.01 | 1197.31 | 19.79 | 80.21 | 4.70 |
| **HC-1** | MZL | 15.16 | 2446.20 | 52.62 | 47.38 | 3.67 |
| **HAIRM** | MZL | 16.74 | 2026.53 | 52.86 | 47.14 | 4.13 |
| **SSK41-Copa** | MZL | 17.24 | 1560.28 | 35.94 | 64.06 | 4.66 |
| **ESKOL** | MZL | 18.18 | 2292.31 | 61.51 | 38.49 | 3.69 |
| **SSK41** | MZL | 18.24 | 1276.55 | 23.21 | 76.79 | 4.74 |
| **SSK41-Vene** | MZL | 23.30 | 1672.02 | 17.32 | 82.68 | 4.95 |
| **VL51** | MZL | 26.42 | 769.82 | 22.06 | 77.94 | 4.09 |
| **Karpas1718** | MZL | 27.01 | 1324.54 | 41.83 | 58.17 | 3.25 |
| **OCILY7** | DLBCL-GCB | 30.79 | 632.08 | 33.92 | 66.08 | 2.76 |
| **JEKO1** | MCL | 32.86 | 358.38 | 27.00 | 73.00 | 4.02 |
| **VL51-Ibru** | MZL | 35.18 | 639.48 | 29.96 | 70.04 | 3.76 |
| **Karpas1718-Ide** | MZL | 37.37 | 713.58 | 54.10 | 45.90 | 3.19 |
| **REC1** | MCL | 43.38 | 849.96 | 44.63 | 55.37 | 4.82 |
| **OCILY10** | DLBCL-ABC | 80.40 | 511.73 | 66.00 | 34.00 | 4.17 |

Legend:

**Sensitivity**

- Less sensitive
- Sensitive

**Internalization**

- High
- Low

**Gene expression**

- High
- Low

**Supplementary Figure S5. Effects of CIGB-552 on apoptosis and cell cycle in MZL cell lines.** (A) MZL cells were treated with CIGB-552 for 72 hours at twice the IC50 concentration. Apoptosis and necrosis were measured using flow cytometry with Annexin-V (FITC) and PI staining, respectively. (B) Cells were exposed to 50 µM of CIGB-552 for 24 hours. Cell cycle analysis was performed via flow cytometry with PI staining. Cell lines are listed according to their associated mechanism of cell death. Representative results from two independent experiments are shown.

**A**

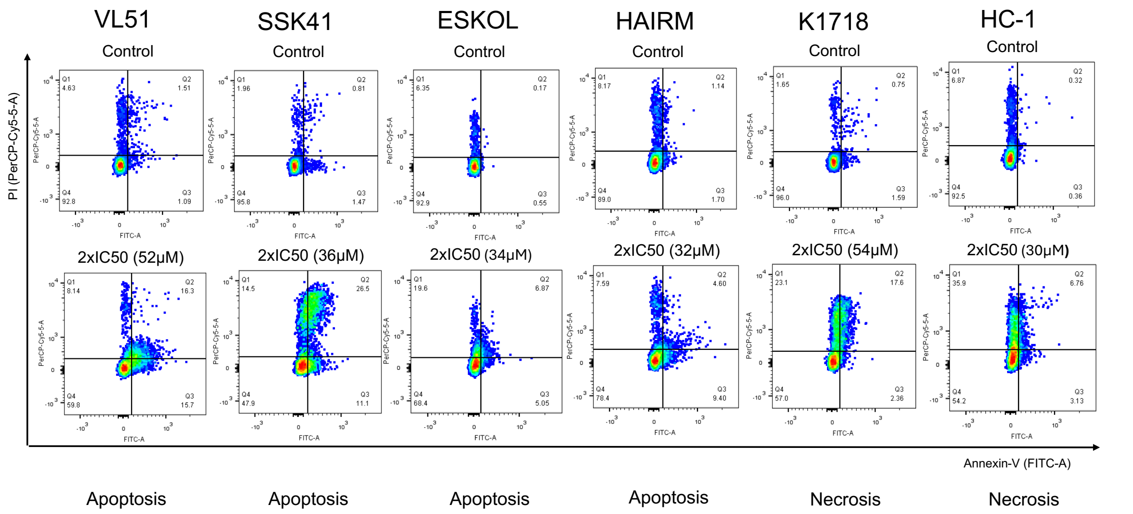


**B**

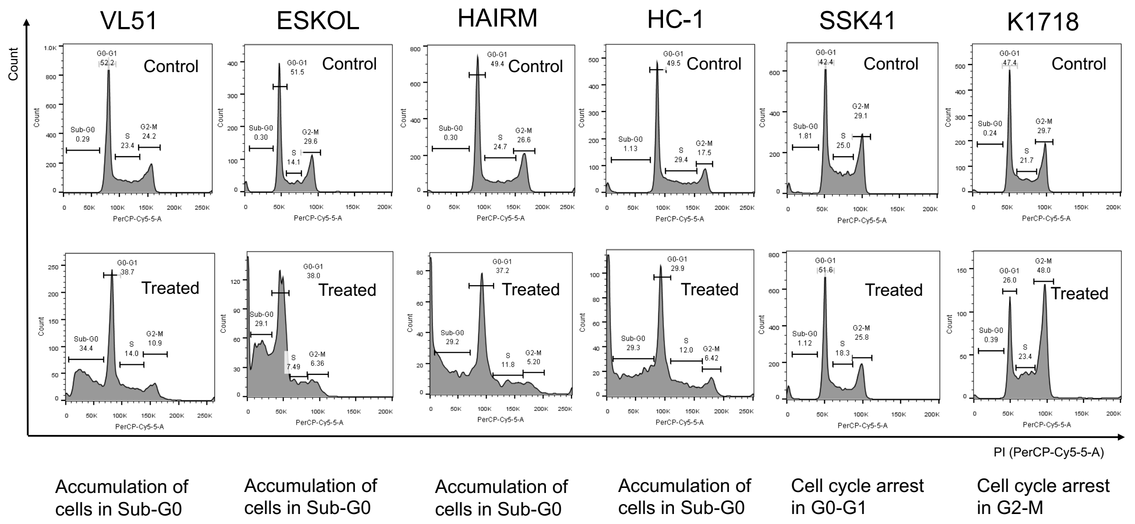

**Supplementary Figure S6. Cell cycle analysis in MZL cell lines at 48h.** VL51, K1718 and SSK41 were exposed to CIGB-552 for 48h at twice the IC50 values. Cell cycle was evaluated by flow cytometry upon PI staining. Representative results of two independent experiments.

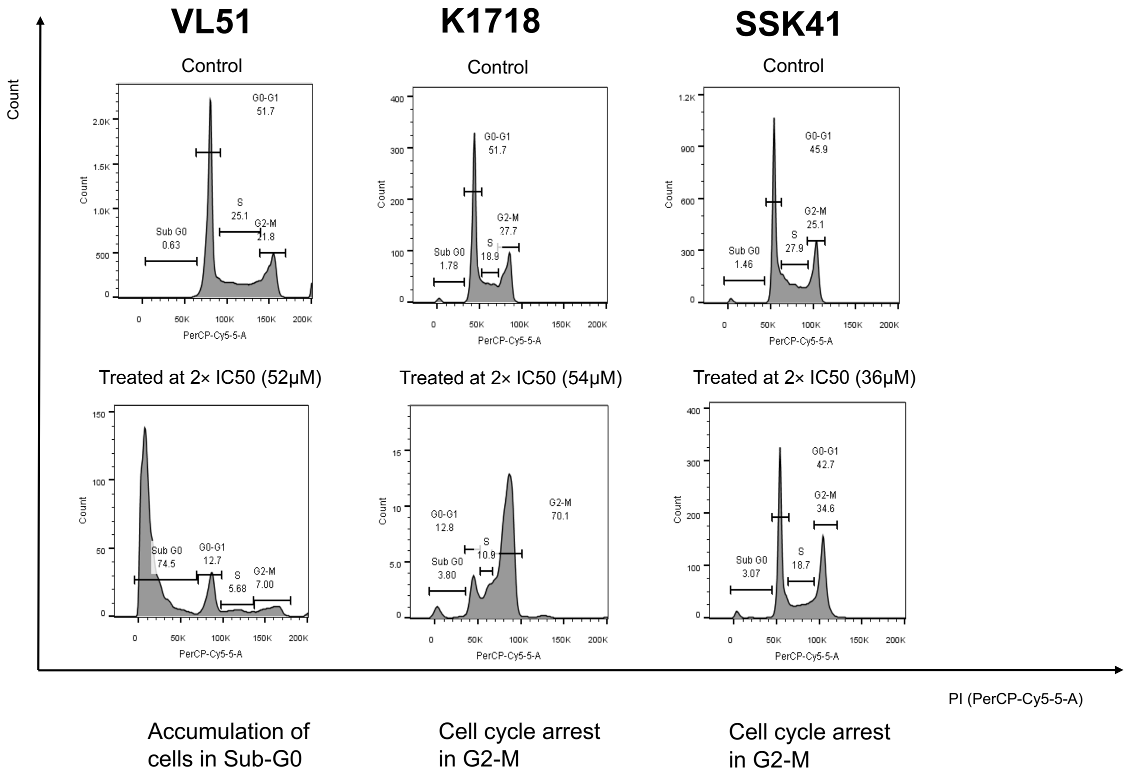

**Supplementary Figure S7. Cell cycle analysis of MZL cell lines exposed to CIGB-552 with and without vitamin E (α-tocopherol, 100 µM) pretreatment.** Karpas1718 and SSK41 cells were treated with 50 µM CIGB-552 for 48h. Cell cycle distribution was analyzed by flow cytometry (PI staining).

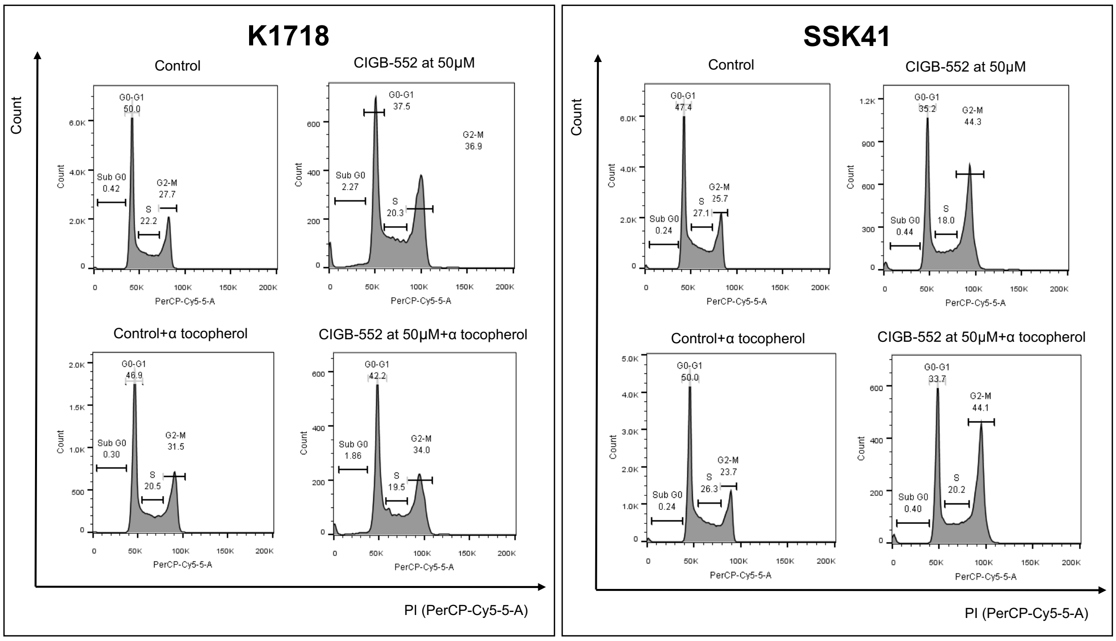

**Supplementary Figure S8. Annexin-V (FITC) and PI staining in MCL (GRANTA-519) and DLBCL-GCB cell lines (OCI-LY7, PFEIFFER, and SU-DHL-16) exposed to CIGB-552 or DMSO**. Cells were exposed to CIGB-552 for 48h (A) and 72h (B) at 50 μM or 120 μM concentration. Apoptosis and necrosis were evaluated by flow cytometry using Annexin V (FITC) and PI staining, respectively.

**A**

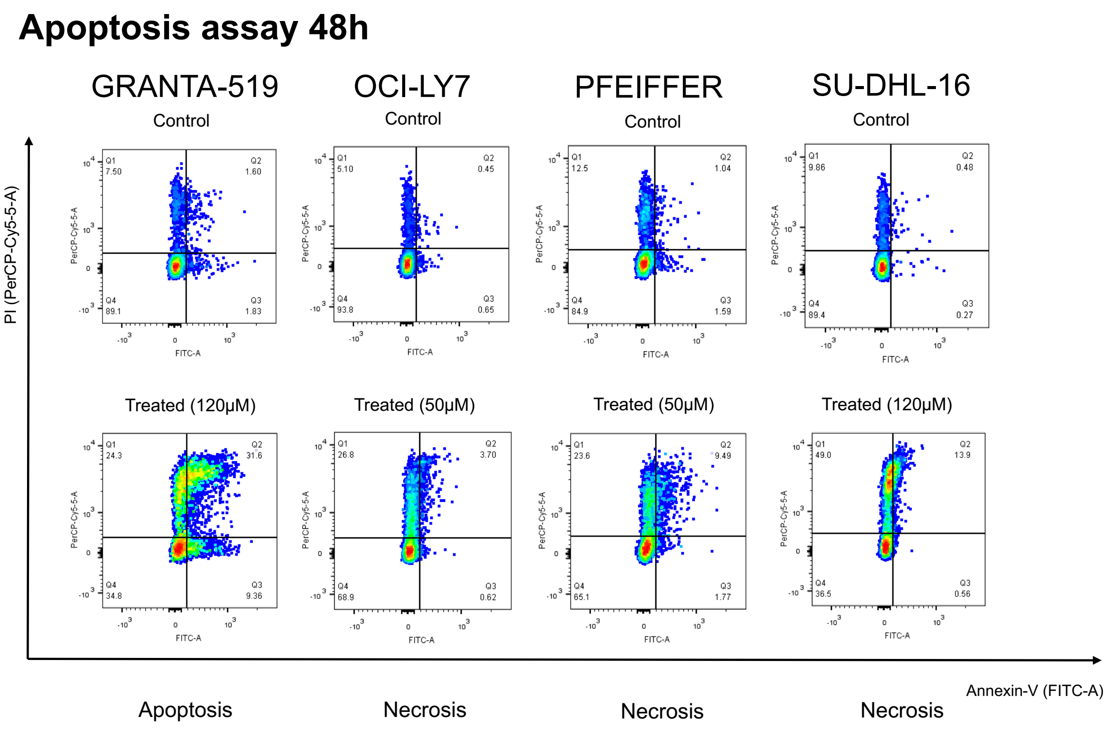


**B**

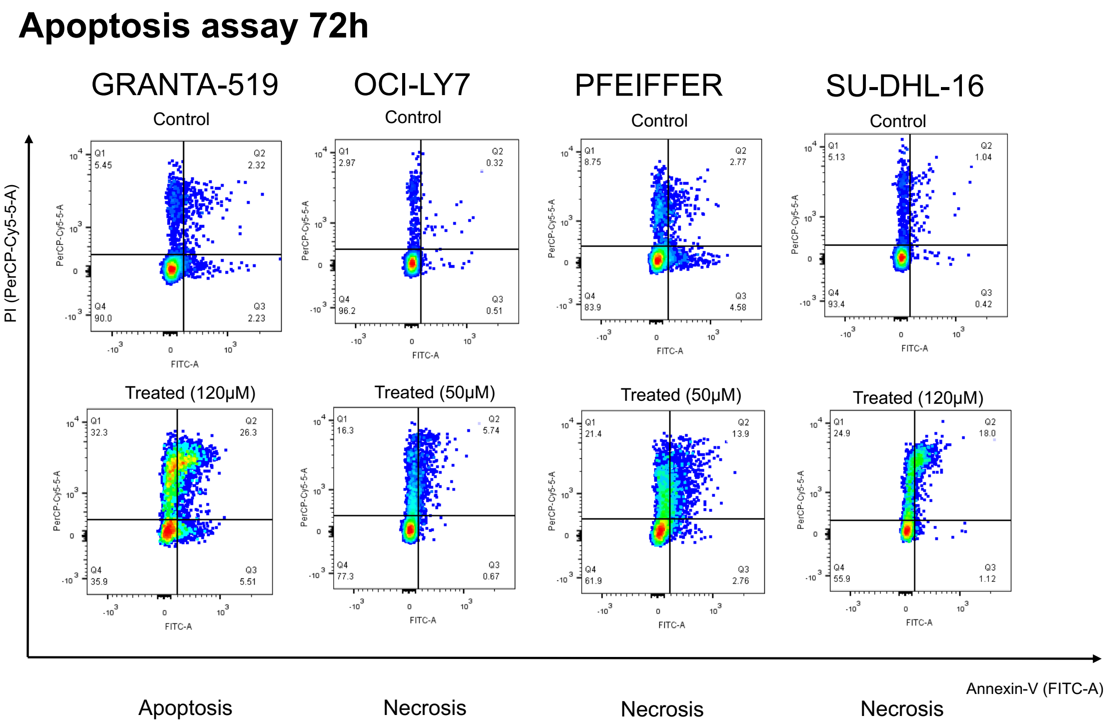

**Supplementary Figure S9. COMMD1 expression in B-cell lymphoma after exposure to CIGB-552.** Immunoblot analysis of COMMD1 protein levels in VL51 cells following 24 h of exposure to CIGB-552 at the IC50 and 2xIC50 values. Vinculin was used as a housekeeping gene.

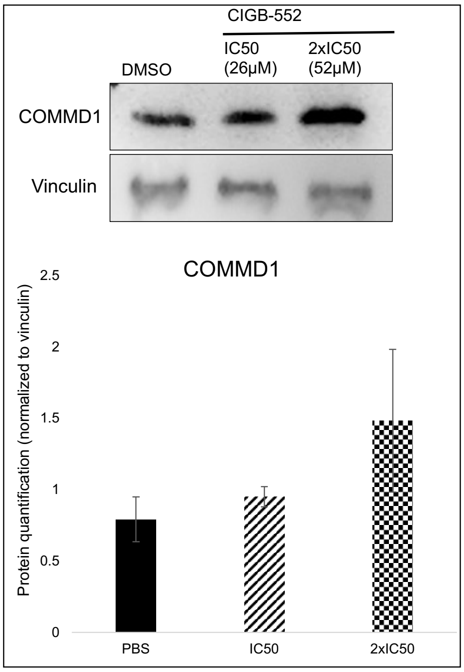

**Supplementary Figure S10. Immunoblotting in MZL cell lines exposed to CIGB-552 or DMSO.** (A) Representative immunoblots and (B) quantification using ImageJ of phospho-RelA (p-RelA, Ser536), total RelA, phospho-AKT (p-AKT, Ser473), total AKT, MCL1, BCL2, and PARP-1 in VL51, K1718, and SSK41 cells following 4 or 24 h of exposure to CIGB-552 at the IC50 and 2× IC50 concentrations. DMSO-treated cells were used as controls, and vinculin served as a housekeeping gene. Statistical significance was assessed using one-tailed paired *t*-tests for all proteins except PARP-1, for which a two-tailed paired *t*-test was used. (C) Representative immunoblots showing phospho-AKT (p-AKT, Ser473), total AKT, phospho-ERK (p-ERK, Tyr204), total ERK, NF-κB1, NF-κB2, and Cyclin D1 expression in VL51 cells following 24 h of exposure to CIGB-552 at the IC50 concentration. Suspension cells were treated in horizontally positioned flasks to maximize the cell culture surface area and facilitate peptide–cell interactions. Vinculin served as a loading control.

**A**

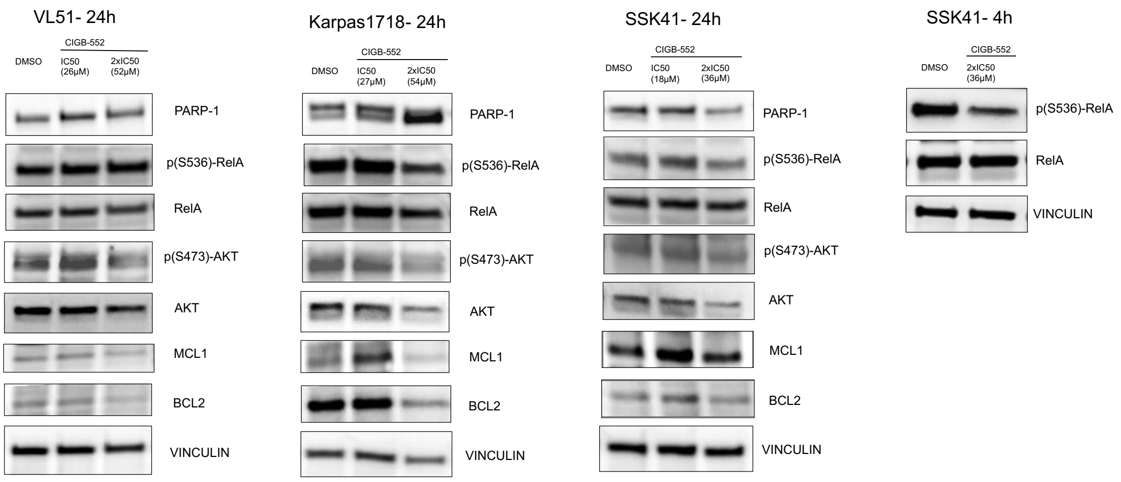


**B**

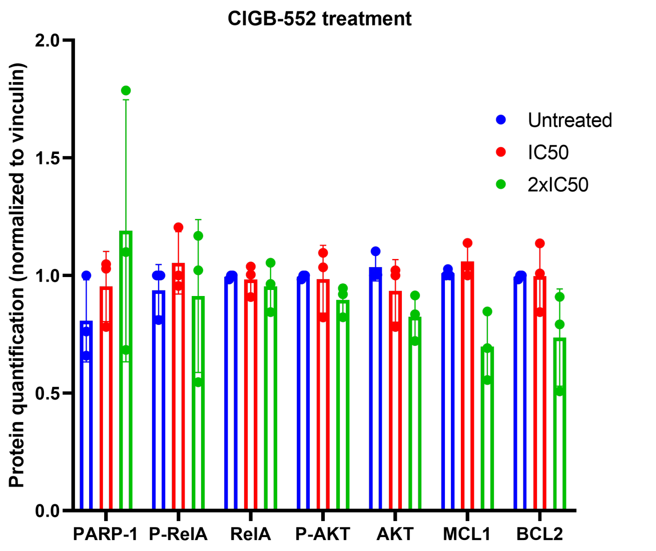


| | proteins | PARP-1 | P-RelA | RelA | P-AKT | AKT | MCL1 | BCL2 |
|---|---|---|---|---|---|---|---|---|
| p-value | IC50 | 0.15178 | 0.142696 | 0.405576 | 0.456189 | 0.143755 | 0.202862 | 0.491334 |
| | 2xIC50 | 0.455917 | 0.463204 | 0.289253 | 0.045366 | 0.012448 | 0.033538 | 0.082923 |

**C**

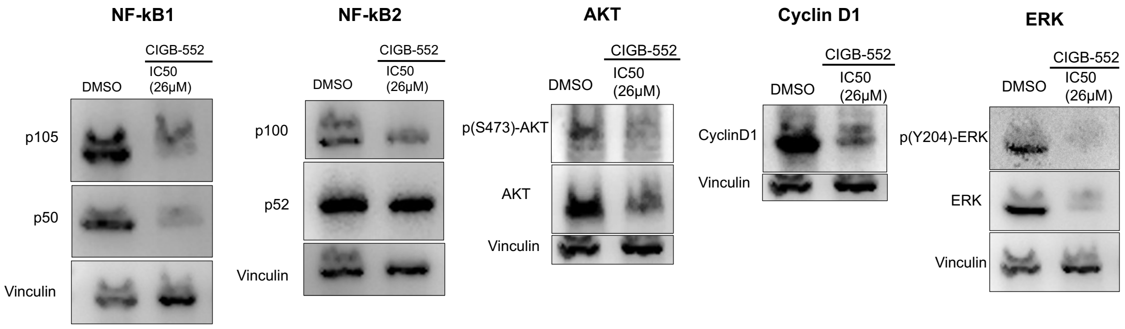


**D**

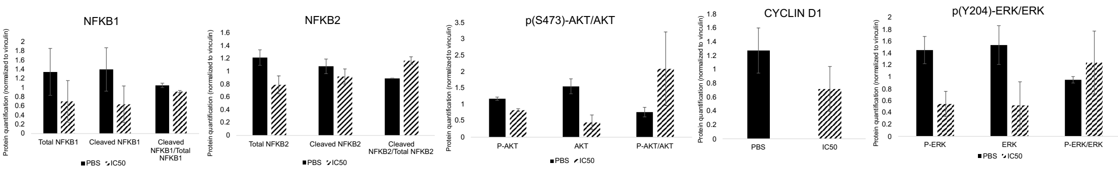


| proteins | NFKB1 total | NFKB1 cleaved | NFKB1 cleaved/total | NFKB2 total | NFKB2 cleaved | NFKB2 cleaved/total | P-AKT | AKT | P-AKT/AKT | CYCLIN D1 | P-ERK | ERK | P-ERK/ERK |
|---|---|---|---|---|---|---|---|---|---|---|---|---|---|
| condition | IC50 | IC50 | IC50 | IC50 | IC50 | IC50 | IC50 | IC50 | IC50 | IC50 | IC50 | IC50 | IC50 |
| p-value | 0.26 | 0.22 | 0.12 | 0.13 | 0.25 | 0.04 | 0.06 | 0.09 | 0.19 | 0.22 | 0.11 | 0.15 | 0.30 |

**Supplementary Figure S11. Combination of CIGB-552 with ibrutinib in MZL cell lines.** Sensitivity to the addition of CIGB-552 to ibrutinib was evaluated by MTT assay at 72h in VL51 (PAR and IBR-Res) in the combination of Ibrutinib 1µM (dil 1:10) + CIGB-552 25µM (dil 1:2), and in K1718 (PAR and IDE-Res), and SSK41 (PAR and VEN-Res) cell lines in the combination of Ibrutinib 100nM (dil 1:3) + CIGB-552 25µM (dil 1:2). DMSO was used as a control. Representative curves of at least two independent experiments.

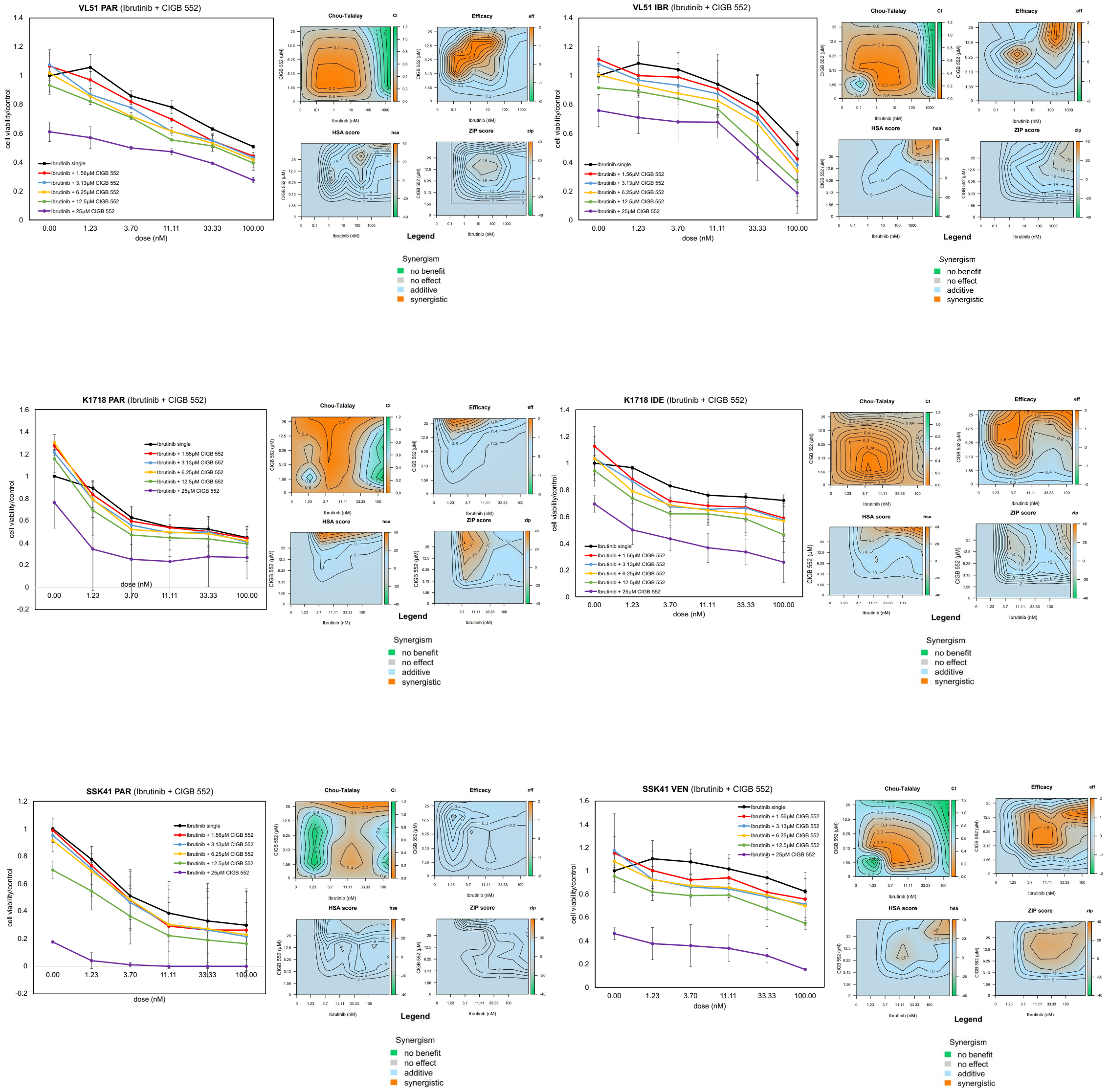

**Supplementary Figure S12. Combination of CIGB-552 with BTK-targeting agents.** Sensitivity to the addition of CIGB-552 to BTK-targeting agents (zanubrutinib, pirtobrutinib and NX-5948) was evaluated by MTT assay at 72h on SSK41 cell line. DMSO was used as a control. Representative curves of at least two independent experiments.

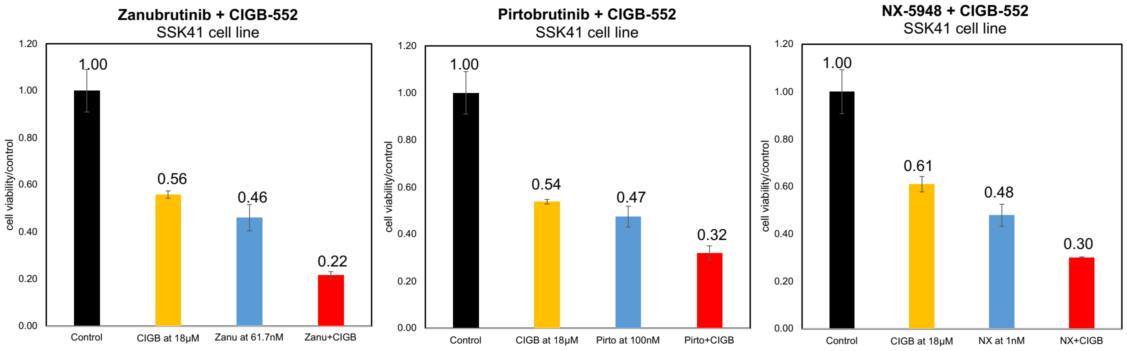

**Supplementary Figure S13. Combination of CIGB-552 with ibrutinib in a canine DLBCL cell line.** (A) The sensitivity to the addition of CIGB-552 to ibrutinib was evaluated by Incucyte assay (6 days of exposure, data normalized at 2d8h0m), combining 0.3 and 1.5 μM of CIGB-552 with 5 pM of ibrutinib. (B) The benefit of the combination was also confirmed by MTT assay at 72h. DMSO was used as a control. Representative curves of two independent experiments.

**A**

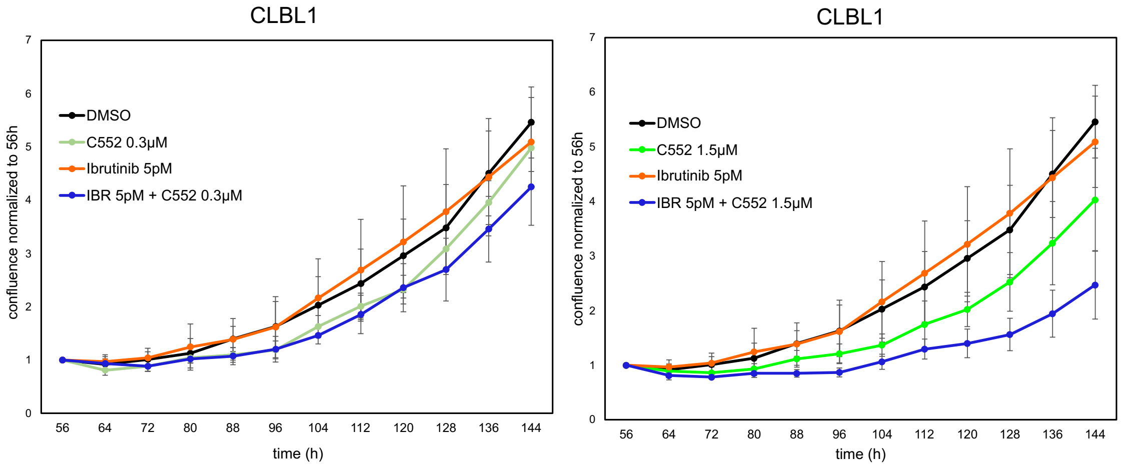


**B**

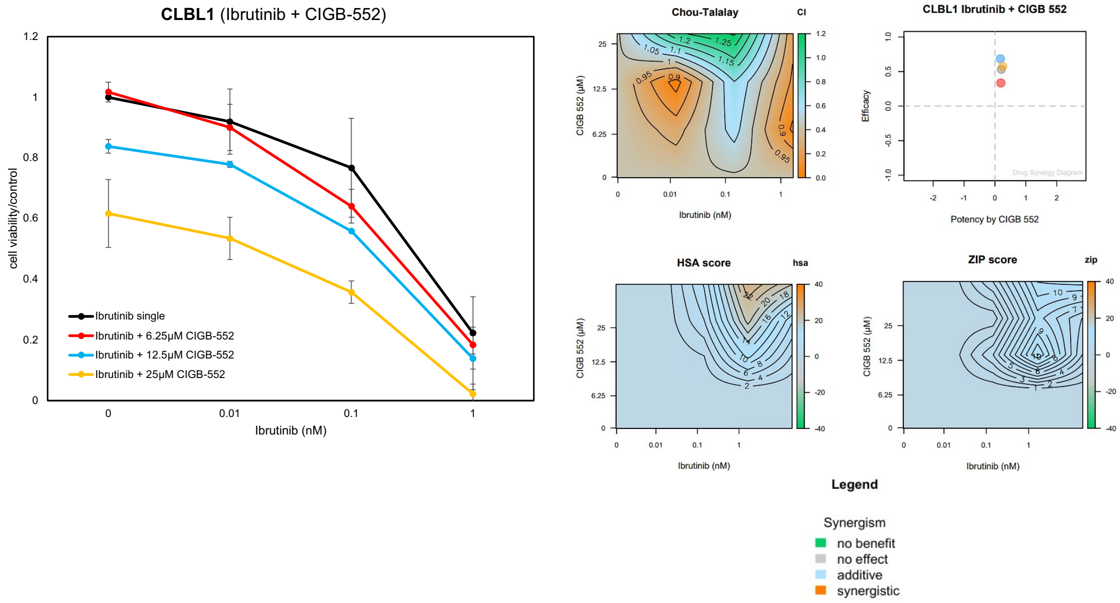
